# Assortative mate choice and epistatic mating-trait architecture induce complex movement of the crow hybrid zone

**DOI:** 10.1101/2020.03.10.985333

**Authors:** Dirk Metzler, Ulrich Knief, Joshua V. Peñalba, Jochen B. W. Wolf

## Abstract

Hybrid zones provide a window into the evolutionary processes governing species divergence. While the role of postzygotic isolation has been extensively characterized in the context of hybrid zones, the contribution of prezygotic isolation is less well explored. Here, we investigate the effects of assortative mate choice, the underlying preference function and mating-trait architecture, and the strength of sexual selection on hybrid zone dynamics. We explore this question by means of a mathematical model parameterized with phenotype and genotype data from the hybrid zone between all-black carrion and grey-coated hooded crows. The best-fit model resulted in narrow clines for two mating-trait loci coding for colour phenotype maintained by a moderate degree of assortative mating. Epistasis between the two loci induced hybrid-zone movement in favor of alleles conveying dark plumage followed by a shift in the opposite direction favouring grey-coated phenotypes ∼1,200 generations after secondary contact. Unlinked neutral loci diffused near-unimpeded across the zone. These results were generally robust to the choice of matching rule (self-referencing or parental imprinting) and effects of genetic drift. Overall, this study illustrates under which conditions assortative mating can maintain steep clines in mating-trait loci without generalizing to genome-wide reproductive isolation. It further emphasizes the importance of mating-trait architecture for spatio–temporal hybrid-zone dynamics.

## 1 Introduction

In sexually reproducing organisms, speciation requires the emergence of reproductive isolation (Coyne and Orr, 2004; Dobzhansky, 1937; Mayr, 1942). Isolating barriers can arise from any mechanism reducing the probability of heterospecific mate choice, fertilization success or hybrid fitness (Coyne and Orr, 2004; De Queiroz, 2007). During the first steps of population divergence, a limited number of loci subject to disruptive or divergent selection can lead to incomplete reproductive isolation (Wu, 2001). Under conditions of gene flow, effective migration will consequently be reduced around these loci (now functioning as “barrier loci”) leading to islands of increased genetic differentiation (Ravinet et al., 2017; Wolf and Ellegren, 2017). Depending on the magnitude of the total selection coefficient relative to recombination, the localized effect around barrier loci may, or may not, generalize and reduce gene flow across the entire genome (coupling) (Barton, 1983; Bierne et al., 2011; Feder et al., 2014). Understanding how traits function as isolating barriers and how they are encoded in the genomes of diverging populations is a crucial step to fully appreciate the mechanisms generating global biodiversity (Dobzhansky, 1937; Presgraves, 2010; Shaw and Mullen, 2011; Wolf et al., 2010).

The above-mentioned processes do not take place in a void, but unfold in populations subject to the governance of space and time (Abbott et al., 2013). During periods of geographic isolation, reproductive barriers most readily evolve as a by-product of divergence and are put to test when populations come into secondary contact (Barton and Hewitt, 1985). When reproductive isolation is complete, sister species can coexist and potentially overlap in their ranges, provided there is little competition. Incomplete isolation will result in hybrid zones (Bigelow, 1965; Harrison, 1990; Hewitt, 1988; Rieseberg et al., 1999). These zones act as semipermeable barriers retaining loci under divergent selection, while unlinked neutral variants can still introgress (Key, 1968; Payseur, 2010). When the homogenizing effect of dispersal into the zone is offset by selection against hybrids, the resulting hybrid zone can reach relative stability in the form of sigmoid clines (“tension-zone model”; Barton and Hewitt, 1985; McEntee et al., 2018). As a result, unique clines are expected for different traits and the underlying genes, unless coupling elicits a genome-wide response (Barton, 1983; Flaxman et al., 2014; Feder et al., 2014). Cline width and position will be influenced by the strength of selection, dispersal distances and the genetic architecture of the trait subject to selection (Endler, 1977; May et al., 1975; Mallet, 1986). Importantly, hybrid zones need not be static. Asymmetries in relative fitness between parental populations or a dominant genetic architecture of the loci contributing to divergence may affect cline centers and induce hybrid-zone movement until they are caught in a trough of low population density, often aligning with physical or ecological barriers (Brodin et al., 2013; Mallet, 1986; Secondi et al., 2006).

Mathematical models and computer simulations within the framework of cline analyses have led to insights on the effects of selection strength (Barton and Hewitt, 1985), the underlying nature of selection (exogenous vs. endogenous) (Kruuk et al., 1999), the genetic architecture of the contributing loci (Bürger, 2017; Mallet and Barton, 1989; Mallet, 1986) and their complex interactions (Barton, 1983; Barton and Shpak, 2000a; Kruuk et al., 1999). Most models, however, rely on the assumption of random mating, and few have explored the role of isolation arising from mate choice alone (but see Irwin, 2020). This gap deserves attention, given the importance attributed to pre-mating isolation, in particular during the early stages of speciation (Coyne and Orr, 1997; Stelkens et al., 2010; Mayr, 1963). Under certain conditions sexual selection can drive rapid evolution of mating cues and preferences promoting speciation (Mendelson et al., 2014), in particular in conjunction with ecological factors (Ritchie, 2007; Servedio and Boughman, 2017) in the form of sensory drive (Seehausen et al., 2008) or magic traits (Servedio et al., 2011).

Yet, one of the major challenges for speciation to occur by sexual section is recombination breaking down the association between mating cues and corresponding preferences (Verzijden et al., 2012; Kopp et al., 2018). This obstacle can be overcome in two ways: physical linkage of the loci underlying genetically determined trait and preference (Merrill et al., 2019; Xu and Shaw, 2019), or a matching rule aligning the phenotype of a genetically encoded mating trait with a corresponding preference. Among the mechanism that can lead to such a matching rule are imprinting on parental phenotypes and self-referencing, that is, individuals inspect themselves and prefer mating partners of a similar phenotype (Hauber et al., 2000; Hauber and Sherman, 2001, 2003; Kopp et al., 2018). Evidence for imprinting of mating traits has been reported for many taxonomic groups, including birds, mammals, fish, amphibians and even insects (see Verzijden et al., 2012, and Kopp et al., 2018, for references). Hauber and Sherman (2001) review evidence for self-referencing in several avian and mammalian species, as for example found in cowbirds (Hauber et al., 2000, see also Hare et al., 2003, and Hauber and Sherman, 2003).

With assortative mating according to matching rules, frequencies of preference and trait are *per se* correlated, which can result in positive frequency-dependent selection. With some degree of incipient divergence (secondary sympatry, geographic isolation) the correlation between trait and preference has the potential to accelerate speciation (Servedio, 2016). Indeed, the preference for sexually selected traits, e.g. as an effect of self-referencing or imprinting on parental phenotypes, has been shown to promote phenotypic divergence and maintain polymorphism in the face of gene flow (Yang et al., 2019; Brodin and Haas, 2006; Verzijden et al., 2012). Empirical work in hybrid zones regularly identifies sexually selected mating traits and preferences as an important component of reproductive isolation (Yang et al., 2019; Schumer et al., 2017; Seehausen et al., 2008; Hench et al., 2019). However, simulation studies suggest that assortative mating can maintain diverged mating traits, but impede genome-wide gene flow of unlinked genetic variation just slightly (Brodin and Haas, 2009; Irwin, 2020).

Here, we investigate the interplay between phenotypic divergence in mating traits, its genetic basis and mating preferences for the well-studied hybrid zone between all-black carrion and grey-coated hooded crows (*Corvus (corone) corone* and *C. (c*.*) cornix*) that presumably arose by secondary contact in the early Holocene approximately 12,000 years (or 2,000 crow generations) ago (Mayr, 1942; Meise, 1928; Parkin et al., 2003; Vijay et al., 2016; Kopp et al., 2018). In this system, there is only limited evidence for post-zygotic, natural selection against hybrids (Saino, 1990; Saino and Bolzern, 1992; Saino and Villa, 1992), but multiple support for plumage-based assortative mating (Meise, 1928; Randler, 2007a) and social marginalization of minority phenotypes (Saino and Scatizzi, 1991; Londei, 2013).

The narrow morphological cline between the two taxa is thus believed to be mainly driven by prezygotic isolation mediated by assortative mating based on cues encoded in plumage pigmentation patterns (Brodin et al., 2013; Kryukov and Blinov, 1994; Meise, 1928; Vijay et al., 2016). Within the same (sub)species, both sexes display the same pigmentation phenotype, suggesting the possibility of mutual mate choice based on plumage characteristics. Plumage pigmentation patterns are encoded by two epistatically interacting, autosomal pigmentation genes (**Fig. 1**) that are subject to divergent selection (Knief et al., 2019). The rest of the genome, however, appears to admix without much resistance (Knief et al., 2019; Poelstra et al., 2014, 2015; Vijay et al., 2016). An ecological contribution to the hybrid-zone dynamics exceeding local effects can almost certainly be excluded (Haas et al., 2010; Randler, 2007b; Rolando and Laiolo, 1994; Saino et al., 1998). The exact mechanism underlying assortative mate choice remains elusive for the crows system. In this manuscript, we focus on parental imprinting or self-referent phenotype matching which have been suggested for crows (Brodin and Haas, 2006; Londei, 2013), and constitute widespread, generic learning mechanisms readily available in birds (Verzijden et al., 2012). The contribution of preference genes in close linkage to the pigmentation loci as e.g. suggested for *Heliconius* (Merrill et al., 2019) remains highly speculative in the crow system (Poelstra et al., 2014) and will not be explored here.

**Figure 1:**
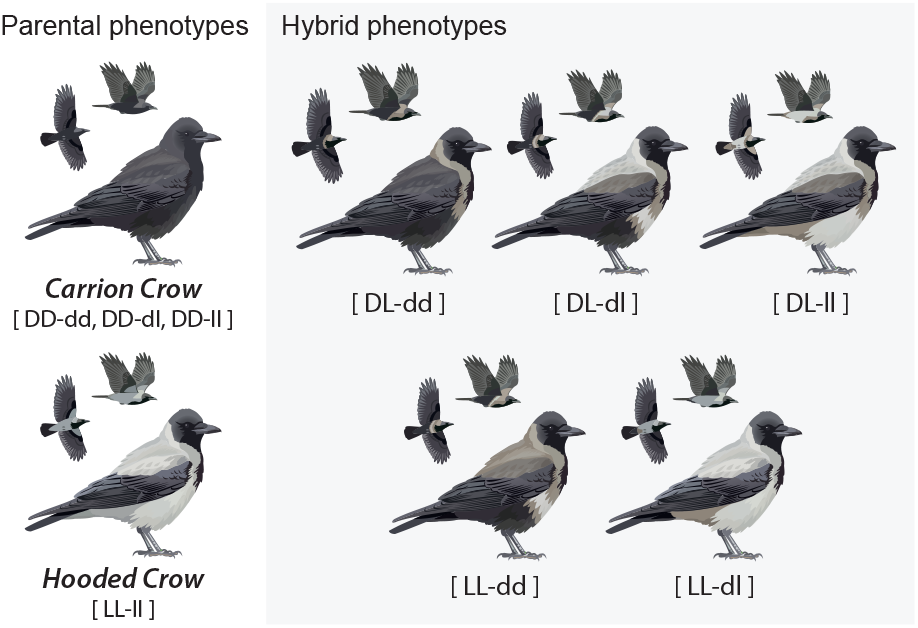
Reconstruction of crow plumage phenotypes as a function of an epistatic genetic architecture of two mating-trait loci as described in Knief et al. (2019). Individuals homozygous for the dark allele (*D*) at locus 1 are always black. The phenotype of heterozygotes (*DL*) and homozygotes carrying the light allele (*LL*) are modified by the genotype of locus 2 (dark: *d* or light: *l*). Only double homozygotes for the light alleles (*LL-ll*) show the pure hooded phenotype. This epistatic architecture underlies our main model. An additive genetic architecture with nine distinct phenotypes is explored separately (2.3.2). Crow images courtesy of Dan Zetterström, modified by Joshua Peñalba.

Here, we formulate a mathematical model invoking assortative mating as a potential driver maintaining phenotypic divergence and fit it to empirical data from the crow system. This merger of mathematical modelling with empirical data reduces the arbitrary parameter space underlying stand-alone simulations. Yet, it still allows exploring central features of general relevance within the confines of a well-investigated empirical system. It thus constitutes a complementary approach to generic simulation studies exploring the role of assortative mating on hybrid-zone dynamics and speciation (Irwin, 2020). Our first aim is to investigate whether essential features of the observed hybrid zone and the available genetic data can be explained by assortative mating according to a matching rule alone without the need to invoke natural selection. We then explore model variants in which central model components are varied: the matching rule underlying mate choice, the genetic architecture of the mating trait and sexual-selection fitness costs of mate choice. Finally, we study whether the models predict speciation for the crow system. Overall, our analyses provide detailed insight into the processes associated with assortative mating and their impact on spatial hybrid-zone dynamics.

## 2 Material and methods

In the following, we will first detail the empirical data and then specify the assumptions and setup of the model. We fit the dispersal parameters (see 2.2.4 below) to data from Siefke (1994), the genotype–phenotype map to data from Knief et al. (2019) collected on adult crows in southern Europe (see 2.1.2 below), and use genetic data from nestlings sampled in the central-European hybrid zone to fit all other parameters of our model, including those for the mating preferences; see online supplement Fig. S1 for an overview.

### 2.1 Empirical data

#### 2.1.1 Phenotypic variation

Details on quantification of phenotypic variation can be found in Knief et al. (2019). In brief, the authors sampled crow individuals across the German and Italian part of the European crow hybrid zone (hereafter “central” and “southern” transects, respectively). Transects were chosen such that they were perpendicular to the geographical course of the zone and included phenotypically pure carrion and hooded crow populations *(Corvus (corone) corone* and *C. (c*.*) cornix*) at either end, as well as hybrids within the hybrid zone. In the southern transect, individuals were sampled as fully feathered adults allowing characterization of individual phenotypes. For a total of 129 individuals, 11 plumage patches were scored for the amount of eumelanin deposited in the feathers (coded as 0, 1, 2 with increasing blackness) and subsequently combined using principal component analysis. Principal component 1 (PC1) explained 78.22 % of the phenotypic variance with positive values reflecting a black carrion-crow phenotype, negative values a grey hooded-crow phenotype and intermediate values being associated with phenotypic variation in hybrids (Knief et al., 2019). In this study, we used these PC1 scores as the metric for a one-dimensional representation of phenotypic variation.

#### 2.1.2 The genetic basis of phenotypic variation

Using genome wide association mapping, Knief et al. (2019) identified two major-effect loci explaining 87.91 % of the variation in plumage patterns represented by PC1. Most of the variation in PC1 (up to 76.07 %) was explained by a locus on chromosome 18 (*chr18*) embedded in a region of reduced recombination. For the purpose of this study, we represent allelic variation at this locus with capital letters as *D* = dark and *L* = light. The second locus, which is associated with allelic variation of the *NDP* gene on chromosome 1, will be denoted in small letters with alleles *d* = dark and *l* = light. Knief et al. (2019) demonstrated that phenotypic variation was best explained by recessive epistasis between *chr18* and *NDP* explaining an additional 10.87 % of phenotypic variation captured by PC1: Allelic variation in *NDP* had no phenotypic effect in all-black *chr18*_*DD*_ individuals, but accounted for most of the residual variation in *chr18*_*DL*_ and *chr18*_*LL*_ (**Fig. 1** and online supplement Fig. S2). Under this model of epistasis, the possible nine genotypic combinations (*DDdd, DDdl, DDll, DLdd, DLdl, DLll, LLdd, LLdl, LLll*) thus reduce to seven phenotypic states, as *DDdd, DDdl* and *DDll* all code for the same black carrion-crow phenotype. **Fig. 1** illustrates the phenotypic space using representative individuals with the respective genotypic constitution.

To explore the effect of genetic architecture on hybrid-zone dynamics in the model (see below), we will consider both the epistatic genetic architecture described above, and phenotypes from an additive model (see online supplement Fig. S2).

#### 2.1.3 Sample preparation and genotyping

To quantify assortative mate choice and predict its effect on the genotypic constitution across the hybrid zone, one would ideally sample breeding pairs along a transect. This is unfortunately not feasible at a larger scale in crows. Instead, we sampled nestlings from the central transect at an age at which plumage characteristics can not be inferred with certainty (Blotzheim et al., 1993); see online supplement Fig. S1. The sampling scheme thus excludes any form of postzygotic isolation after the nestling stage. Due to the relatedness of nestlings, their genotypes contain information on the genotypic constitution of their parents and hence on phenotype-dependent mate choice. We therefore used the genotypic distribution of 152 nestlings sampled from 55 nests (median brood size = 3 nestlings) along the transect (Knief et al., 2019) as read-out for model fitting (see 2.2.6). This procedure rests on the assumption that the genetic architecture of plumage patterning is equivalent in both transects. This is well supported by a shared evolutionary history of the parental populations at either side of the hybrid zone (Vijay et al., 2016), congruent landscapes of genetic variation across the central and southern part of the hybrid zone (Vijay et al., 2016) and near-identical selection signatures at the underlying loci (Knief et al., 2019).

We genotyped all individuals from the central transect for a selection of 1,152 SNPs spread across the whole genome using the GoldenGate assay (Illumina). A detailed description of the assay design, sample preparation, SNP calling and quality control procedure is given in Knief et al. (2019). The final data set comprised all 152 individuals genotyped at 1,111 polymorphic loci (average call rate of 99.48%). We further included 65 individuals of the allopatric populations in the GoldenGate genotyping and added five hooded crows that had been sequenced on the HiSeq2000 (Illumina) platform (paired-end libraries; sequence coverage ranged from 7.12 × to 13.28 ×, average = 9.77 ×, median = 9.83 ×) and genotyped using the HaplotypeCaller in GATK (v3.3.0; DePristo et al., 2011; Vijay et al., 2016). In those five individuals, we found 104 inconsistencies out of 5,501 genotypes (1.89%). All individuals were sexed based on their heterozygosity for 114 SNPs located on the sex chromosome Z (excluding the pseudo-autosomal region located at chrZ ≤ 2.56 Mb, N = 15 SNPs). Genotypes of the pigmentation loci (see 2.1.2) were inferred as follows. The genetic factor on *chr18* was represented by ancestry-informative diplotypes spanning the 2.8 Mb region on chromosome 18. Ancestry (pure carrion *chr18*_*DD*_; pure hooded *chr18*_*LL*_; mixed *chr18*_*DL*_) was inferred using the NewHybrids software (v2.0+ Developmental. July/August 2007; Anderson and Thompson, 2002) as described in Knief et al. (2019). Individuals that were assigned as backcrosses with evidence for recombination were excluded (*N* = 12 individuals in the hybrid zone and *N* = 11 allopatric individuals). To represent genotypic variation in *NDP*, we chose the SNP on chromosome 1 at 6,195,380 Mb (genome version 2.5, RefSeq Assembly ID according to the National Center for Biotechnology Information: GCF 000738735.1; Poelstra et al., 2014) that explained most of the variation in plumage coloration (Knief et al., 2019) and is linked to the likely causal structural mutation (Weissensteiner et al., 2020).

#### 2.1.4 Paternity assignment

Our model assumes that individuals from the same nest are full-siblings. Thus, we estimated whether nest mates were full- or half-siblings originating from extra-pair copulations using 752 SNPs that were located on all autosomes except chromosome 18 (for details see Knief et al., 2020). Full- and half-sibling pairs could reliably be separated based on their kinship coefficients (*θ*, Weir et al., 2006). For one nest containing three individuals, with two full-siblings (*θ* = 0.23 and *θ* = 0.26) and one half-sibling (*θ* = 0.12), relationships could not unambiguously be resolved. We removed this nest (N = 3 individuals) and another six extra-pair young, leaving 52 nests containing a total of 131 individuals in the hybrid zone and 64 allopatric individuals for subsequent analyses.

### 2.2 Theoretical model

Here we specify our main model for the hybrid zone of carrion crows and hooded crows. The model parameters were then fit to data from the crow system (sections 2.2.4 and 2.2.6). For an overview of the different components of our models and the data sets to which we fit them, see online supplement Fig. S1. In section 2.3, we will specify three variants of our model, which differ from the main model in their assumptions on mating preferences (2.3.1), the genetic architecture of the mating trait (2.3.2) or the fitness costs associated with mate choice (2.3.3). The source code of the programs that we developed to carry out simulation studies and to fit all four model variants to the data is available from https://github.com/statgenlmu/assortative_crows/.

#### 2.2.1 Temporal dimension

Similar to other European taxa, Eurasian carrion and hooded crows were presumably separated into different refugia during the last ice age (Mayr, 1942; Hewitt, 1988) which peaked in glacial coverage around 20,000–18,000 years ago (Hewitt, 1988; Frenzel, 1992). Despite a subsequent period of increasing temperature, it was likely only after intermission of a cooling phase in the younger Dryas 12,800 to 11,500 years ago (Goslar et al., 2000; Rasmussen et al., 2014) that rapid warming opened up for suitable crow habitat including breeding opportunities in larger shrubs or trees (Giesecke et al., 2017). We therefore set the initiation of secondary contact to approximately 12,000 years before present. Using a generation time of 6 years for the Eurasian crow (Kutschera et al., 2020; Vijay et al., 2016), this corresponds to 2,000 generations.

#### 2.2.2 Spatial dimension

In our model, we conceptualize space in a one-dimensional grid of 200 bins. When we fit the model to the crow data, we assume that these bins represent 5-km-wide strips that are parallel to the initial contact line (ICL) of the two color morphs, such that our model covers a range from 500 km to the west to 500 km to the east of the ICL. More precisely, we assume that bins 100 and 101 represent the areas that are directly adjacent to the ICL, up to 5 km to the west and east, respectively; bins 99 and 102 are the areas that are 5 to 10 km western or eastern of the ICL, etc. Generally, bin *x* for any *x* ∈ {1, 2, …, 200} represents the area that has a distance between |*x* − 100|·5 km and |*x* − 101|·5 km to the ICL and is west of the ICL if *x* ≤ 100 or to the east for *x* ≥ 101. The precise north–south extension of these strips in not important for our model as long as it is long enough to avoid boundary effects in our dispersal model (section 2.2.4 below).

#### 2.2.3 Modeling approach for assortative mating

We assume that each bin is populated by the same number of crows and that this number is so large that we can neglect genetic drift and any other source of random fluctuations in genotype frequencies, which means that our model is deterministic. For each genotype *g* with respect to the two mating-trait loci, let *f*_*x,g*_(*t*) ∈ [0, 1] be the frequency of *g* in bin *x* at time *t*. We assume discrete time steps *s* = 0, 1, 2, … corresponding to non-overlapping generations, with time *t* = 0 referring to the time of the secondary contact of carrion crows and hooded crows after the ice age. We assume that at time 0 all crows in bins 1 to 100 were carrion crows and all crows in bins 101 to 200 were hooded crows. For *x ≥* 101 this implies *f*_*x,g*_(0) = 1 if *g* = *LLll*, and *f*_*x,g*_(0) = 0 otherwise. For *x ≤* 100 we obtain *f*_*x,g*_(0) = 0 for *g* ∉ {*DDdd, DDdl, DDll*}, and assume Hardy–Weinberg frequencies (*f*_*x,DDdd*_ (0), *f*_*x,DDdl*_(0), *f*_*x,DDll*_(0)) = *δ*^2^, 2 · *δ* · (1 − *δ*), (1 − *δ*)^2^, where *δ* is the initial frequency of allele *d* in bin *x*, which we assume to be equal in all bins *x* ≤ 100 at time 0.

The core of our model is the recursion to compute the generation *t* + 1 genotype distribution matrix *f* (*t* + 1) = (*f*_*x,g*_(*t* + 1)) _*x,g*_ from the genotype distribution matrix *f* (*t*). Implicitly, we assume that the sex ratio is the same for all bins and all genotypes. Following a common mass-action law approach (see e.g. Otto and Day, 2011) we assume that the number of matings between individuals with genotypes *g*_1_ and *g*_2_ taking place in bin *x* is proportional to the number of potential couples in bin *x*, weighted by a mate choice value 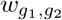 (with 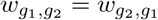 ; see 2.2.5 below). The latter can be interpreted as the probability that a male and a female crow with genotypes *g*_1_ and *g*_2_ will mate if they meet. Note that in this setup it is not relevant whether the female, the male or both sexes are taking the mating decision based on the similarity of their phenotypes.

If *d*_*z,x*_ is the dispersal probability from bin *z* to bin *x* (section 2.2.4), the number of matings between individuals of genotypes *g*_1_ and *g*_2_ taking place in bin *x* is proportional to

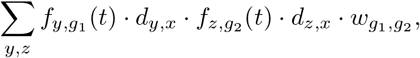

where the summation variables *y* and *z* iterate over all combinations of bins from *x* − 10 (or 1 if *x* − 10 ≤ 0) to *x* + 10 (or 200 if *x* + 10 > 200), with the range of ± 10 bins reflecting a maximum dispersal distance of 50 km (section 2.2.4). This proportionality could arise, for example, if within each bin randomly moving individuals form a pair at the beginning of the mating season and occupy one of a limited number of territories to reproduce, as is observed in the field by Blotzheim et al. (1993). Individuals that need to search longer for suitable mating partners have smaller chances of finding a vacant spot for their nest. Thus, a form of sexual selection emerges even though we assume monogamy. In accordance with this heuristic consideration, our model implies that individuals that are surrounded by fewer preferred mating partners will on average have fewer offspring. Thus, assortative mating induces a reduction in fitness via frequency-dependent selection in both sexes.

To finally obtain the recursion to compute *f* (*t* + 1), let for each triplet (*g, g*_1_, *g*_2_) of genotypes be *µ*(*g* |*g*_1_, *g*_2_) the probability that an offspring of parents with genotypes *g*_1_ and *g*_2_ has genotype *g*, according to Mendelian genetics with free recombination between the two loci. The frequency of genotype *g* in nestlings of generation *t* + 1 in bin *x* is then

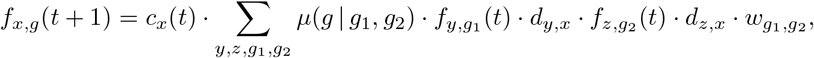

where the value of *c*_*x*_(*t*) ensures that ∑_*g*_*f*_*x,g*_ (*t* + 1) = 1.

#### 2.2.4 Dispersal

Natal dispersal distance was assumed to be equal for both sexes and parameterized with data from Siefke (1994, Table 3 and table caption), who analyzed data of hooded and carrion crow individuals that were ringed as nestlings and recovered during the breeding period. Kalchreuter (1970) investigated ring recovery data of carrion crows and obtained a similar distribution of dispersal distances. To use the dispersal distances quantified by Siefke (1994) on a two-dimensional map, we first needed to project these distances to our one-dimensional representation of the contact zone. For this we assume that the ICL is a straight line going in south–north direction and decompose each dispersal movement of a crow into its component *D*_*x*_ orthogonal to the ICL and its component *D*_*y*_ parallel to the ICL. The values of *D*_*x*_ or *D*_*y*_ are positive, if the direction of movement is west to east or south to north, respectively, and negative otherwise. We assume that the random distribution of the dispersal vector (*D*_*x*_, *D*_*y*_) is a mixture of two spherical two-dimensional normal distributions. A single spherical normal distribution did not lead to a satisfying fit to the long-tailed distribution of the dispersal distance data (online supplement Fig. S4 top). Thus, the dispersal vector can be represented as (*D*_*x*_, *D*_*y*_) = *S* · (*Z*_*x*_, *Z*_*y*_), where *S* is a random variable with two possible values *σ*_1_ and *σ*_2_ and the vector (*Z*_*x*_, *Z*_*y*_) has a two-dimensional standard normal distribution. To fit the three parameters *σ*_1_, *σ*_2_ and *p* = Pr(*S* = *σ*_1_) to the empirical dispersal distances (Siefke, 1994) we numerically maximized the likelihood of (*σ*_1_, *σ*_2_, *p*) using the method of Byrd et al. (1995) as implemented in the R command `optim` with the option `L-BFGS-B` (R Core Team, 2018). To compute likelihoods we used that with probability *p* the squared rescaled dispersal distance ‖ (*D*_*x*_, *D*_*y*_)*/σ*_1_ ‖^2^ or otherwise ‖ (*D*_*x*_, *D*_*y*_)*/σ*_2_ ‖^2^ is equal to 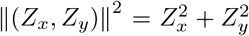 and thus chi-square distributed with 2 degrees of freedom. The fitted values are 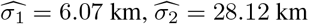 and 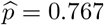.

As the bins in our models represent areas that extend as long stripes parallel to the ICL, only the component *D*_*x*_ is relevant. Thus, for the probability that a crow stemming from bin *x* is found in bin *x* + *d*, the dispersal distance *d* is a discretization of *D*_*x*_, whose distribution consists of 76.7 % of a normal distribution with a standard deviation of 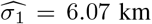, with the other 23.3 % coming from a normal distribution with a standard deviation of 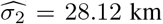. For computational efficiency we restrict the range of *d* to ±10 bins, corresponding to a max. ±50km, setting Pr(*d* = − 10) = Pr(*d* = 10) = Pr(*D*_*x*_ *<* − 47.5) = Pr(*D*_*x*_ > 47.5) (online supplement Fig. S4 bottom). Furthermore, the boundaries of our binned space are reflecting. That is, probability weights of dispersal *d* that would result in *x* + *d <* 1 or *x* + *d* > 200 are shifted to instead increase the probabilities of bin 1 + |*x* + *d*|or 200 − |*x* + *d* − 201|, respectively.

#### 2.2.5 Mate choice

In the absence of decisive knowledge on the mechanism underlying mate choice in the crow system, and for computational feasibility, we assume self-referent phenotype matching in our main model. Using alternative models (see sections 2.3.4, 3.5 and 3.6) we further explore whether this simple matching rule can serve as a proxy for maternal or paternal imprinting (Verzijden et al., 2005, 2012), as has also been suggested for the crow system (Brodin and Haas, 2006, 2009; Londei, 2013).

We summarize the phenotype by the values of the first principal component derived in Knief et al. (2019) (see 2.1.1 and its encoding by two-locus genotypes as specified in section 2.1.2). For simplicity, phenotype values are scaled to a range between 0 (pure hooded crow phenotype, genotype *LLll*) and 1 (pure carrion crow phenotype black, genotype *DDdd, DDdl, DDll*).

Let *ϕ* be the genotype–phenotype map. The mate choice value 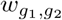 then is the mating probability of two birds with phenotype values *ϕ*(*g*_1_) and *ϕ*(*g*_2_) relative to the mating probability of two birds of equal color, such that 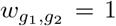. We model the mate choice function *w* with two parameters *η* and *ζ*, where *η* is the mating preference of crows with very different phenotypes and *ζ* controls the width of a Gaussian kernel modeling how mating preference 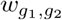 of more similar birds depends on their phenotypic difference *ϕ*(*g*_1_) − *ϕ*(*g*_2_):

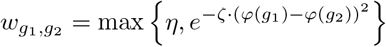

Online supplement Fig. S5 shows how 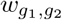 depends on *ϕ*(*g*_1_) − *ϕ*(*g*_2_), with parameter values *η* and *ζ* set according to maximum-likelihood values in our main model. Examples for resulting mate-choice values for various parameter settings are shown in **Fig. 2 B**.

**Figure 2:**
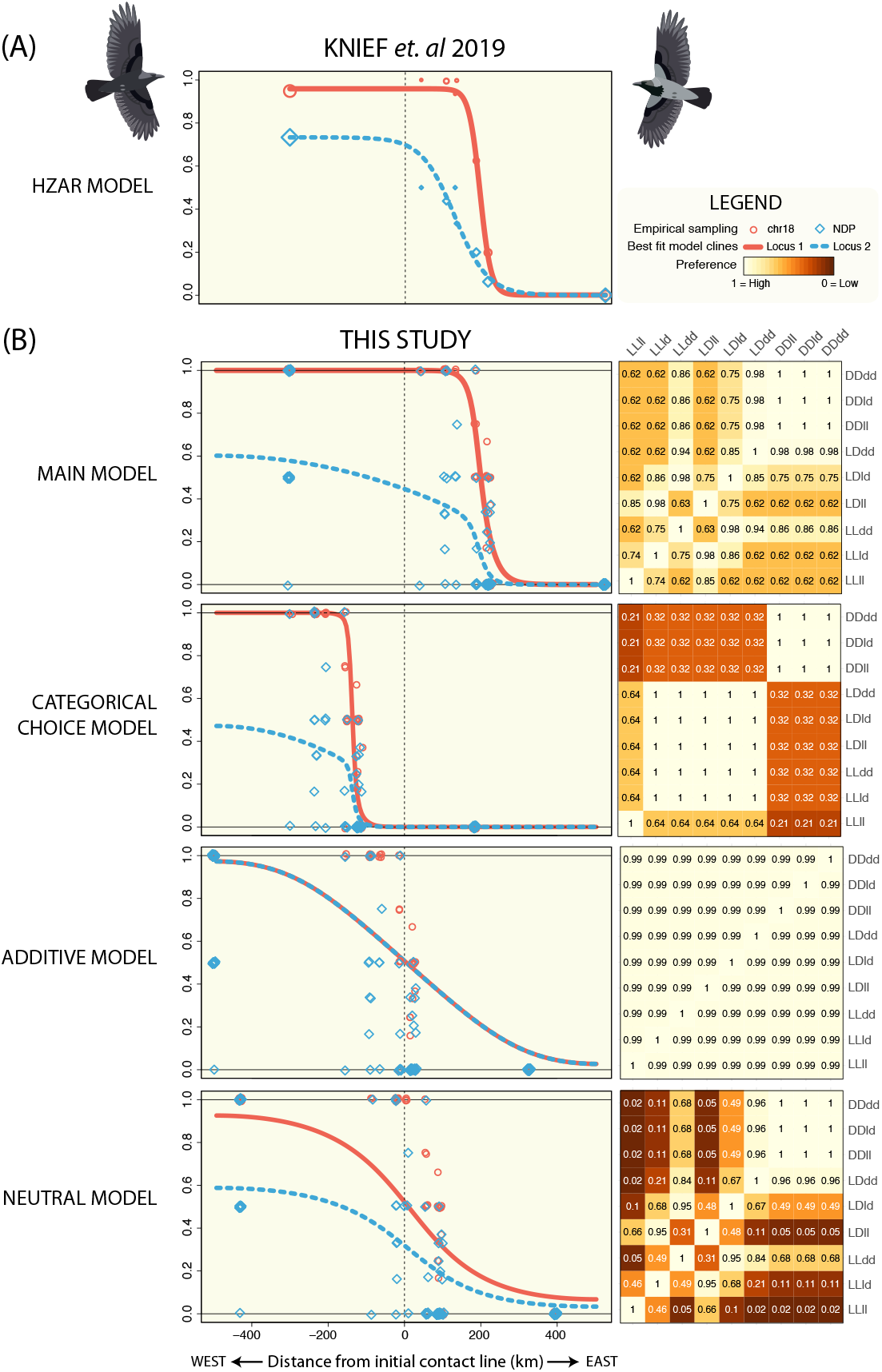
**A:** Clines fitted by Knief et al. (2019) for frequencies of the dark alleles from locus 1 (*chr18* shown in red) and locus 2 (*NDP* shown in blue) of current-day empirical data collected across the German hybrid zone. Allele frequencies were averaged per sampling location and symbol areas reflect sample sizes. The geographic cline models were fitted using the hzar package (version 0.2-5; Derryberry et al., 2014) in R. **B:** Best-fit parameter combinations of the main model and three variants thereof (categorical mate choice, additive genetic architecture, no sexual selection). *left*: Geographic clines of the two mating-trait loci after 2,000 simulated generations shown in relation to the initial zone of secondary contact between carrion crows in the west and hooded crows in the east. Model predictions shown for the dark alleles of locus 1 (red line) and locus 2 (blue line) are overlaid on current day, empirical allele frequencies for each nest: red open bullets and blue diamonds represent the dark allele frequencies at locus 1 and locus 2, respectively. *right*: Matrix *w* of pairwise mate choice values by genotype (mutual mate choice of both sexes); Light yellow represents the highest possible value of 1.0 (no avoidance). Black (which does not occur here) would refer to a value of 0.0, which means that crows of the two genotypes never mate. An intermediate value of e.g. 0.5 reflects half the probability of mating relative to the phenotypic combination with the highest value.

#### 2.2.6 Fitting hybrid-zone model parameter to genotype data

Weissensteiner et al. (2020) inferred that the *d* allele of the *NDP* gene arose approximately 500,000 years ago and segregated in the ancestor of both carrion and hooded crows. Since the three genotypes *DDll, DDld* and *DDdd* all lead to the all-black phenotype, we therefore allow that the ancestral frequency of allele *d* in the west can initially be different from 1. This initial frequency of the *d* allele *δ* is the third model parameter besides *ζ* and *η*. A fourth parameter is the geographic location of the initial contact line. For this, all sampling sites are projected on a one-dimensional transect as specified in Knief et al. (2019), see also section 2.1.3. The position on the transect is measured in kilometers, starting with the most western sampling site as km 0. For the position of the initial contact line (which is assumed to intersect the transect in a right angle) we allow the range from km 300 to km 700 on the transect.

To numerically optimize all four parameters, we first simulated the hybrid-zone model for any triple of candidate values for the first three parameters. Then we calculated for the grid km 300, km 301, …, km 700 the probability of the empirical genotype data for the case that the initial contact line was at that position. To calculate the parameter likelihoods, that is the probability of the data, we rounded the nest locations on the transect to the bins and calculate for each possible combination of parental genotypes the probability that the chick genotypes of a nest stem from such parents (assuming Mendelian inheritance and no segregation distortion as Knief et al., 2020, found no evidence for biased sex ratios in F1-hybrids or for postzygotic isolation). The likelihood calculations were then based on the simulated genotype distribution after 2,000 generations and mate choice according to the model assumptions. Thus, we optimized the likelihood of the four parameters with respect to the genotype data. For this we used the parallel version of the L-BFGS-B method provided by the R (version 3.5) package optimParallel (Gerber and Furrer, 2018).

To infer the point of inflection of a cline from allele frequencies *y*(*x*) in all bins *x* ∈ {1, …, 200} we first chose the bin *x* with the largest value of |*y*(*x* − 1) − *y*(*x* + 1)|, then fitted a third order polynomial to *y* in the range from *x* − 2 to *x* + 2 and calculated the point of inflection of the fitted polynomial.

### 2.3 Model variants

In the following, we formulated deviations from our main model exploring several parameters we deem of importance in the crow system and beyond. These include i) the specific mode of how phenotypic differences are translated into mate choice (preference function), ii) the impact of the genetic architecture of the mating trait and iii) how the system is affected if we remove sexual selection altogether while maintaining assortative mate choice.

#### 2.3.1 Categorical choice model

In this variant of our model, hybrids are summarized into a single category and have no mating preferences. However, carrion crows prefer carrion crows and hooded crows prefer hooded crows as mating partners. If two crows interact and both are of the same category—carrion crow, hooded crow or hybrid—their mating preference value *w* is 1. If one is a hybrid and the other is a hooded crow or a carrion crow, their mating preference value *w* is given by the parameter *ψ*_0_ or *ψ*_1_, respectively. If one is a hooded crow and the other a carrion crow, their mating preference value *w* is the product *ψ*_0_ · *ψ*_1_. We combined this model with our standard genotype–phenotype map as specified for the main model (section 2.1.2).

#### 2.3.2 Additive model

This model variant assumes an additive genetic architecture of plumage pigmentation patterns. The corresponding genotype–phenotype map was combined with the mate-choice function of the main model (section 2.2.5). We constructed the additive genotype–phenotype map by taking the same phenotypic values for the *LLll, LLdl* and *LLdd* genotypes as in the epistatic model and the slope between *LLdl* and *DLdl* to infer all other additive phenotypic values (online supplement Fig. S2). Note that according to this model, only genotype *DDdd* leads to an entirely black carrion crow phenotype. As we assume that shortly after the ice age all crows in the west were black, the initial allele frequency of the *d* allele at locus 2 was 1.0. (See online supplement section G.3; model variant with a relaxation of this assumption.)

#### 2.3.3 Neutral model

Assortative mating according to matching rules usually induces frequency-dependent sexual selection, as finding mating partners may be more costly for rare phenotypes (Kopp et al., 2018; Irwin, 2020). To examine the extent to which sexual selection affects hybrid-zone dynamics, we define a neutral model, in which sexual selection is eliminated by compensating the fitness cost of preferring rare phenotypes as mating partners. Thus, in this variant of our model, all individuals have the same expected number of offspring neglecting the possibility that the increased effort of rare phenotypes to find a mating partner may induce fitness costs. (Factoring out the potential relevance of individual-based recognition and local dominance hierarchies for reproduction in corvids (Chiarati et al., 2010; Holtmann et al., 2019), this assumption seems not too far-fetched.)

Given the narrow width of the hybrid zone, costs for surveying larger distances for this highly mobile species appear minor. Accordingly, in the neutral variant of our model, crows stemming from areas in which their phenotypes are rare will tend to ultimately mate in areas in which the frequency of their matching mating partners is higher. Thus, effective migration patterns are not symmetric and lead to slight differences in population densities that depend on the phenotype. Therefore, to avoid induction of sexual selection by local competition, we have to assume that local population densities can vary keeping only the global population size constant. This may run counter to ecological intuition, but we make this assumption to obtain a neutral model that we can compare to our main model in which selection is induced. As a consequence, the neutral model posits that the total number of individuals in a bin can vary among the bins. We combined this model with the standard genotype–phenotype map of the main model (section 2.1.2).

For the precise specification of this model, let *f*_*x,g*_(*t*) (still) be the frequency of *g* in bin *x* at any time *t* relative to the *initial* total frequency ∑_*g*_ *f*_*x,g*_ (0) in bin *x* (or any other bin), but note that, differently than in the main model, sum ∑_*g*_ *f*_*x,g*_ is in general not 1 for *t ≥* 1. The initial matrix *f* (0) is the same as in 2.2.3. The recursion to calculate *f*(*t*+1) is now

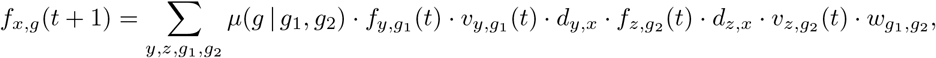

where the factors *v*_*x,g*_(*t*) compensate induced fitness effects for individuals stemming from bin *x* and having genotype *g*, which could reflect, e.g., that these individuals intensify or extend their search for mating partners by a factor of *v*_*x,g*_(*t*), with the assumption that this is possible without incurring any fitness cost. For this, the values of *v*_.,._(*t*) must simultaneously for all (*y, g*_1_) fulfill

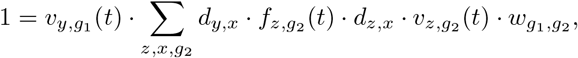

which we solve numerically (see online supplement F for details).

#### 2.3.4 Finite-size population models with self-referencing or imprinting

In the models specified above we neglect random variation such as genetic drift which is clearly present in real populations of finite size. To assess how robust our results are against random fluctuations of this kind we performed simulations using indivdual-based models. In brief, the Central European crow habitat is idealized as a two-dimensional grid of 1000 × 500 territories, each spanning one square kilometer and containing one crow nest (which lies within the large empirically documented range, Blotzheim et al., 1993). These models only consider genetic drift and do not account for other sources of randomness and spatial or temporal variation of ecological conditions. The same assumption on crow dispersal and mating preferences are used as in the main model (see online supplement J for details).

The individual-based model approach not only allows comparing the results from the main model to a finite-population-size model with a self-referent matching rule (online supplement section J.1). Its inherent flexibility also opens the oppportunity to examine the effect of imprinting in which females prefer males that resemble the average phenotype of her parents (online supplement section J.2), males that resemble their fathers or, in another variant of the model, their mothers (online supplement section J.3).

## 3 Results

### 3.1 Main model: parameter estimates

The main model assumes an epistatic genetic architecture of the mating trait, and sexual selection induced by assortative mating favoring common phenotypes. This model produced a good fit to the empirical genotype frequency distribution, and with a log-likelihood of -175.6 was superior to the alternative models discussed below. (Note that these models have the same numbers of parameters.) The cline for locus 1, which is responsible for the majority of the overall phenotypic variation, was steepest (**Fig. 2)**. It was closely followed by a shallower, and geographically offset cline of the second, epistatically interacting locus. This is concordant with the clines that have been fitted to the empirical data in (Knief et al., 2019). At the most western end of the transect, 34 single individuals were sampled from different nest, such that the allele frequencies shown in **Fig. 2 B** in the west are all 0, 0.5 or 1. The local genotype frequencies are shown in online supplement Fig. S3.

Consistent with the proposition that the *light* allele of the second locus already segregated in the ancestral population of carrion crows (Weissensteiner et al., 2020) the maximum-likelihood parameter estimate of the initial allele frequency of the *dark* allele in carrion crows was below one (0.777). As maximum likelihood parameter values for the mate choice function we found (*ζ, η*) = (2.17, 0.617) and kilometer 302 on the transect as position of initial contact. Individuals differing most extremely in appearance (i.e. pure hooded crows with genotype *LLll* vs. pure carrion crows with genotypes *DDdd, DDdl* or *DDll*) were only 61 % as likely to choose each other for reproduction compared to pairs of individuals of equal phenotypes. Note that the ultimate mating rates not only depend on this preference matrix but are also contingent on the frequency of each genotype in the local population determining the relative frequency of mutual encounter.

### 3.2 Hybrid-zone movement

The maximum-likelihood parameterization of the main model predicted a substantial shift of the hybrid-zone center to the east after 2,000 generations. More precisely, the inflection points of loci 1 and 2 were inferred to be 189.6 and 188.3 km to the east of the initial contact line (ICL). To further explore these dynamics we simulated 6,000 generations under the main model that we parameterized with the maximum likelihood estimators described above. During the first ∼ 1,200 generations, the zone moved eastward from the ICL (in favor of dark morphs). After allele *d* of locus 2 decreased to a certain frequency, the hybrid zone began to reverse its movement and shifted westward (now in favor of the light morphs, **Fig. 3** and online supplement I.1).

**Figure 3:**
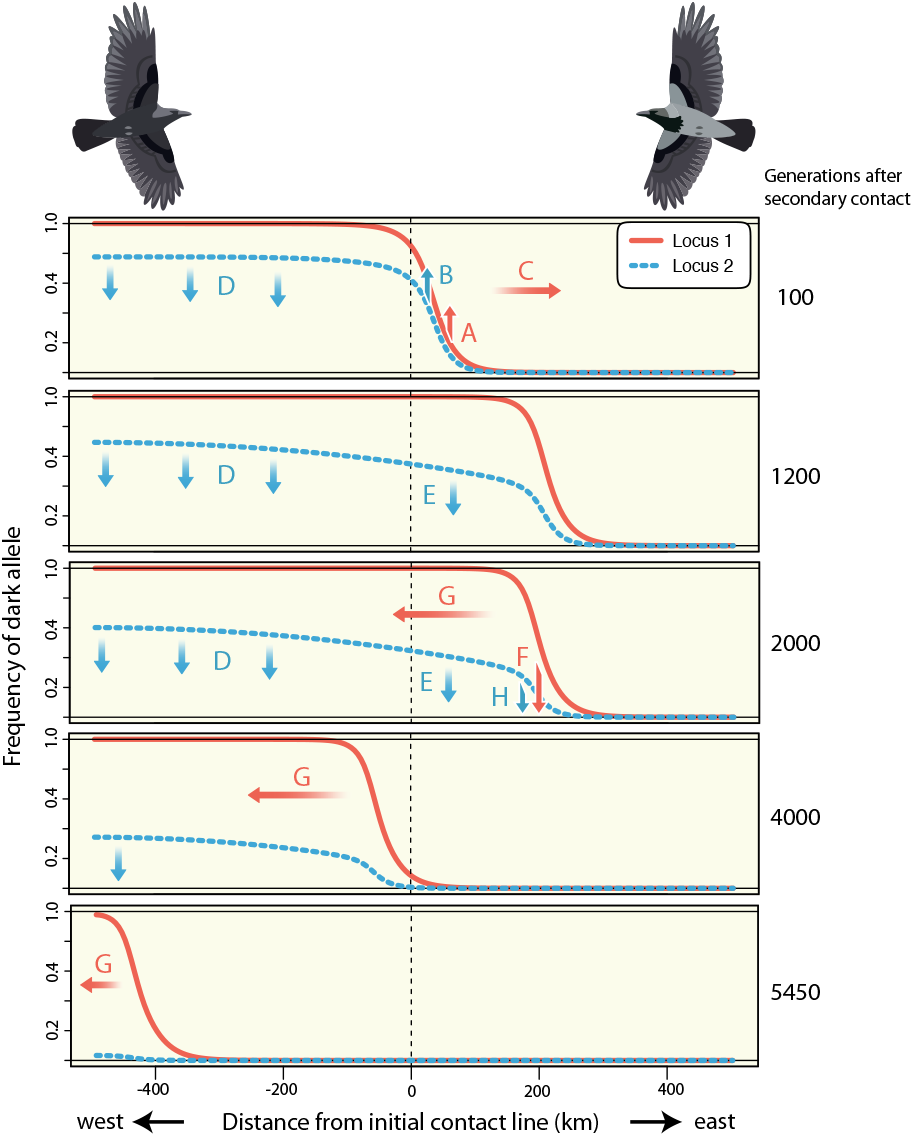
Hybrid zone movement through time shown for both mating-trait loci. As in Fig. 2 model predictions are shown for the dark alleles of locus 1 (red line) and locus 2 (blue line). According to the main model with sexual selection, the hybrid zone first shifts eastward, then reverses its direction westward with carrion crows eventually going extinct. For a detailed description of the arrows accompanied by letters refer to the text.

The movement of the hybrid zone can be explained as follows. Due to the epistatic architecture, three different genotypes code for all-black crows (**Fig. 1**). This induces a high initial frequency of dark morphs among hybrids which induces positive frequency-dependent sexual selection in favor of darker crows. This, in turn, increases the frequency of alleles *D* and *d* in the hybrid zone (**Fig. 3**, arrows A and B) which promotes a shift of the hybrid zone to the east (arrow C). At the same time, the *l* allele moves into the west. While allele *L* is under negative selection in the west, allele *l* has no fitness effects. It freely introgresses into the *DD* background, which entails a decrease of allele *d* (arrow D). In the long run, this also decreases the frequency of *d* in the proximity of the hybrid zone (arrow E), which reduces the fraction of dark morphs among the hybrids and thus the fitness of dark crows. This eventually brings the hybrid-zone movement to a halt after approximately 1,200 generations. As the decrease of allele *d* continues (arrows D and E), sexual selection provides an advantage to lighter morphs, decreasing the frequencies of *D* and *d* in the hybrid zone (arrows F and H). As a consequence, the hybrid zone travels westward (arrow G) and finally drives alleles *d* and *D* to extinction after 5,500 generations.

### 3.3 Model variants

#### 3.3.1 Categorical choice model

With a log-likelihood of -199, the model fit the data substantially worse than the main model assuming mate discrimination between all seven phenotypes. In this model, mate-choice-relevant phenotypes were categorized into pure and hybrid-type. Maximum likelihood estimates for the mating preference parameters *ψ*_0_ and *ψ*_1_ of hooded and carrion crows, respectively, were 0.318 and 0.645, which translates into a larger difference in mate choice **(Fig. 2D)**. As in the main model, pure morphs had a clear preference for phenotypically matched partners. The probability of mating between carrion and hooded crows, however, was only 21 % and thus only about a third of that in the main model (61 %). According to the best fitting model parameter values, also for locus 2 the dark allele *d* was fixed in the west before secondary contact.

In terms of spatio–temporal dynamics according to this model, the hybrid zone had shifted westward since the initial contact. The inferred inflection points of clines of loci 1 and 2 after 2,000 generations were located 142.1 and 141.7 km to the west of the ICL, respectively. According to simulations with more than 2,000 generations, the model predicts a continuous westward shift of the hybrid zone until carrion crows go extinct (online supplement Fig. S15).

#### 3.3.2 Additive model

Changing the genetic architecture from epistasis between the mating-trait loci to additivity had a strong impact. The best possible fit to the data for the additive model was poor (log likelihood -228.7), and maximum likelihood estimates of *ρ* = 2.878 and *η* = 0.987 implied only very slight mating preferences overall **(Fig. 2)**. Mate choice probabilities only differed by 1.3 % (0.987 vs. 1.0) indicating very weak sexual selection. It is thus not surprising that clines of both mating-trait loci were shallow. Still, the difference of 1.3 % in mate-choice probabilities shielded against gene flow, and clines were more pronounced than by simple diffusion of a neutral locus (see section 3.4). Furthermore, due to additivity the frequency curves of the two alleles were still almost identical after 2,000 generations (**Fig. 2)**. With inferred points of inflection of both clines only 0.1 km to the east of the ICL, almost no shift of the clines was predicted. In summary, the additive model could not explain some essential features of the empirical data, highlighting the importance of the genetic architecture of mating traits for hybrid-zone dynamics.

The above results are based on the assumption that, initially, all crows in the west were black, which entails that the dark alleles *D* and *d* were fixed in the west. This differs from the main model, where the frequency of the *d* allele was free to vary. In online supplement section G.3 we report results of further parameter combinations of the additive model relaxing the assumption that all parental carrion crows were entirely black upon secondary contact. Also with this assumption, clines were either not steep enough to fit the empirical data or the empirical disparity in cline location and inclination between the loci was absent. Overall, none of the models with additive genetic effects fit the data well.

#### 3.3.3 Neutral model

Next, we formulated a model without sexual selection where all individuals had the same expected number of offspring and did not incur any fitness advantage or disadvantage in relation to their phenotype. With a log likelihood of -210.3 the fit to the data was substantially worse than in the main model including sexual selection. The best fitting parameter combination for the neutral model had an initial *d* allele frequency for locus 2 of 0.619, and showed a slight shift for both loci 1 and 2 of 11 and 7.2 km to the east from the ICL (measured at the inflection points of the clines).

The maximum-likelihood parameter estimates of the mating function *ζ* = 5.53, *η* = 0.0237 translate to the most extreme avoidance of non-self phenotypes of all investigated model alternatives. Matings of the opposite morph (carrion with hooded crows) were 50 times less likely than between equivalent phenotypes (**Fig. 2)**. Similar as for the categorical model this may be explained by the requirement of maintaining a narrow hybrid zone against increased gene flow in the center of the zone by hybrids traveling far to seek an appropriate partner (with no cost in this model).

While the model somewhat mimicked the empirical disparity in allele frequency clines between loci, it did not capture the steepness of the cline for locus 1. A steeper cline could be achieved by setting the model parameters to values reflecting stronger assortative mate choice. In this case, however, the frequency curve for locus 2 was shifted too far to the east. This would produce many *LLdd* individuals in the hybrid zone, which were near-absent in the empirical data, and explains the poor fit (compare online supplement Fig. S7 to online supplement Fig. S8 and online supplement Fig. S9).

Interestingly, the model predicted a decrease in population density in the center of the hybrid zone accompanied by a population increase at a distance of around 200 km on either side (online supplement Figs. S7 and S9, bottom). Assuming no cost of mate choice and movement, this shift in densities is due to individual compensation for differences in sexual attractiveness induced by the preference function. As we assume that individuals of a phenotype that is rare in their environment intensify their search for mating partners, these individuals may effectively migrate over larger distances and deplete local population densities.

### 3.4 Effects of assortative mating on unlinked neutral loci

Using the best-fit parameters of the main model, we ran additional simulations in which we included a third, neutral locus. This bi-allelic locus was unlinked to the loci associated with the mating trait and had no effect on mate choice or dispersal. We defined the focal allele as the allele with initial frequency of 1 in the west and of 0 in the east. We compared this simulation to a simulation in which all three loci had no effect on the mating trait or dispersal (random mating) and essentially follow neutral diffusion according to our dispersal model (section 2.2.4). (Note that in the latter case, locus 1 and locus 3 have the same allelic distribution.)

The spatial distribution of alleles from the neutral locus 3 was affected by the mating-trait loci, but this effect was very weak (**Fig. 4** and online supplement Fig. S16). After 2,000 generations the frequency of the neutral allele at locus 3 was 0.620 at the western boundary of the range and 0.398 in the far east in the main model with locus 1 and 2 under sexual selection. With random mating, the frequency of locus 3 was slightly closer to equalized allele frequencies (at 0.5) with 0.597 in the west and 0.403 in the east. Hence, this model predicts that sexual selection does not substantially reduce effective migration at loci unlinked to those determining the mating trait. Even if assortative mating in this model entails that hybrids have reduced fitness, local reduction in gene flow of the mating trait affected other parts of the genome only marginally.

**Figure 4:**
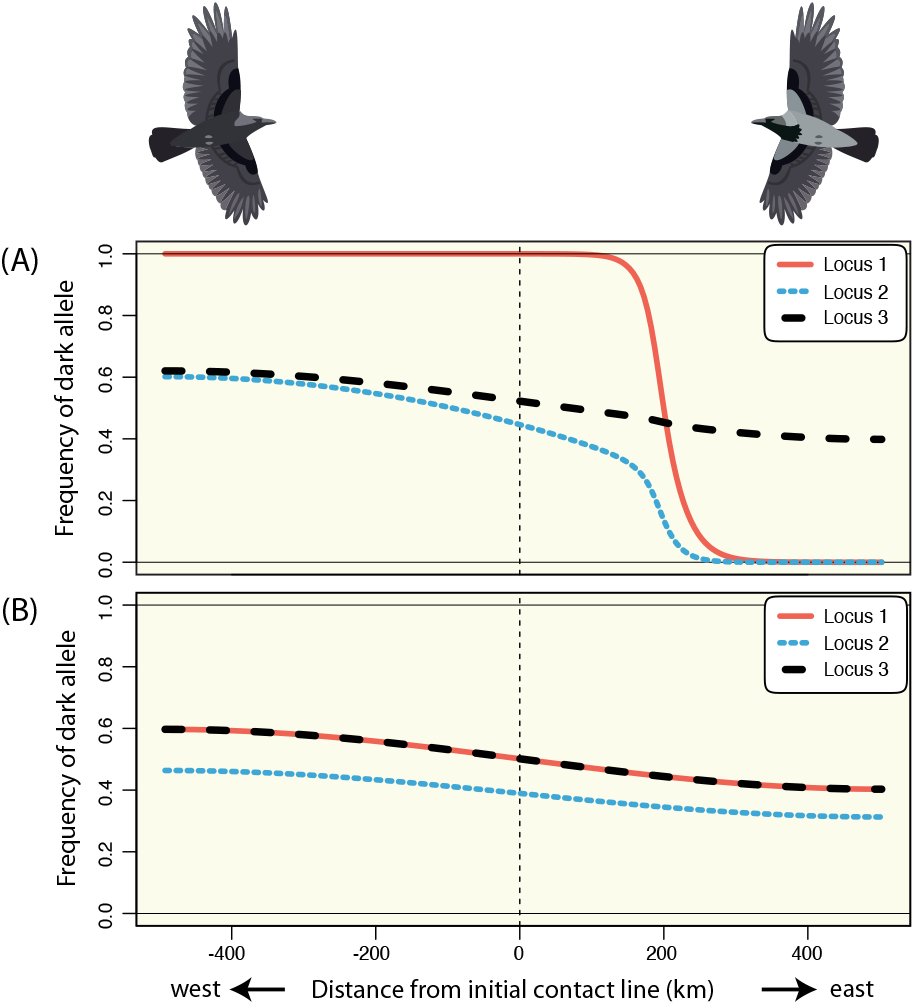
**A:** Clines resulting from the sexual selection model with the best-fitting parameters for locus 1 (red), locus 2 (blue) and a third, neutral locus (black). **B:** Clines for the same three loci resulting from models with the same initial conditions and dispersal as in the sexual selection model, but simulated with random mating.

### 3.5 Effect of genetic drift

The finite-population-size models allow for stochastic variation in the spatial distribution of genotypes (online supplement Fig. S17), and ultimately cline dynamics (online supplement Fig. S20). The finite-population model with the self-referent matching rule, parameterized according to our main model, showed the same type of hybrid-zone cline dynamics as our main model (**Fig. 5 B**; online supplement section J.1 and Fig. S18). Hence, the outcome was not qualitatively altered by genetic drift.

**Figure 5:**
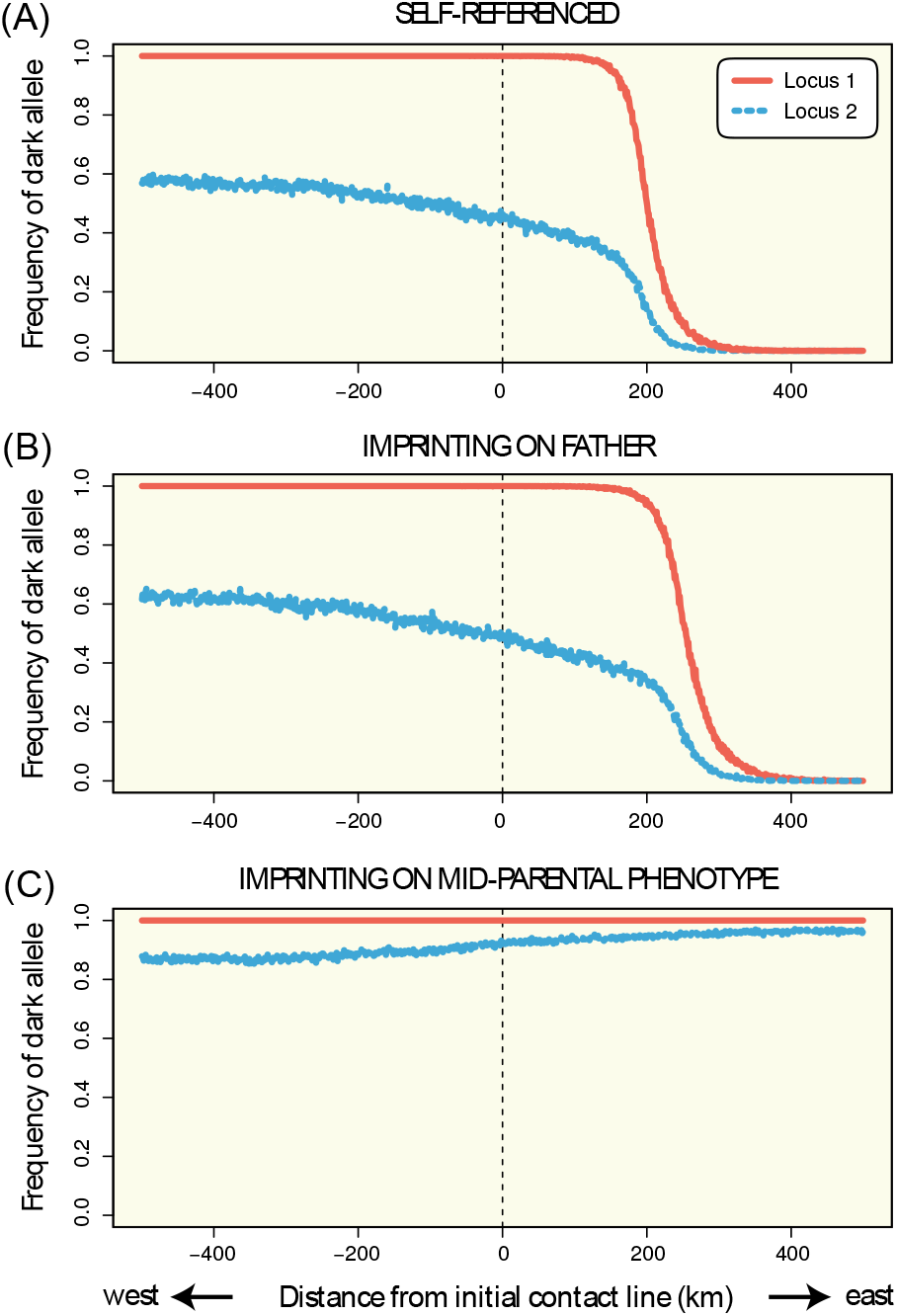
Simulation results for individual-based models, which allow for genetic drift, for the time point after 2000 generations.

### 3.6 Imprinting vs. self-reference

The flexibility of the individual-based models further allowed exploring the effect of the matching rule. Specifically, we assessed whether self-referent phenotype matching would serve as a good proxy for imprinting, as has been suggested before (Verzijden et al., 2012, 2005). Imprinting was modelled as female-based mate preference based on the phenotype of one parent (online supplement section J.3) or on the average phenotype of her parents (online supplement section J.2). In the first case, results closely aligned with the outcome based on self-referencing, regardless which sex served as the imprinting model (**Fig. 5 B** and online supplement Fig. S20). In case of imprinting on the mid-parental phenotype, cline dynamics were substantially different, Here, the hybrid zone moved eastward until the carrion-crow phenotype became fixed (**Fig. 5 C**). Another differences was that the locus-2 cline was shifted slightly to the east compared to the locus-1 cline, such that directly in the hybrid zone after 1000 generation the locus-2 dark allele was more frequent than the locus-1 dark allele (online supplement Fig. S19).

## 4 Discussion

In this study, we explored the effects of prezygotic isolation on hybrid zone dynamics. Specifically, we studied how the interplay of a genetically-determined mating trait and preference for the trait based on simple matching rules (self-referencing, imprinting) affects hybrid zone position and width through time. Analyzing empirical data from the European crow system by simulation-based model-fitting allowed us to estimate the strength and mode of mate choice as a function of individual genotype combinations. Clearly, every theoretical model is a necessary simplification of reality (Popper, 1934; Box, 1979). We chose to neglect ecological factors and did not explicate spatial details of dispersal barriers despite known effects on hybrid zone dynamics (Barton, 1979; Barton and Hewitt, 1989). We rather focused on the question of whether intrinsic behavioral properties of the system suffice to explain the phenotypic and genotypic composition across the crow hybrid zone for which rich data has now been collected for nearly a century (Meise, 1928; Knief et al., 2019). We reach the conclusion that assortative mating based on learning of phenotypic cues alone can explain empirical observations to a great extent under realistic assumptions and model parametrization. For a better understanding of the underlying processes, we examined the effects of the mate-choice function, the genetic architecture of the mating trait, the fitness costs induced by frequency-dependent selection and finite population size using model variations. We discuss these aspects in turn with a focus on their relevance for reproductive isolation. First, we home in on the mating-trait loci themselves and then consider genome-wide implications.

### 4.1 Mating-trait loci

Theory predicts that mating preferences based on matching-rules, such as self-referent phenotype matching or imprinting, readily enhance positive frequency-dependent sexual selection of locally abundant mating-trait alleles (Yeh and Servedio, 2015). In the context of secondary gene flow between populations with diverged mating-trait values, this process is expected to result in directional selection on each side of the hybrid zone pulling in opposite directions. For the entire population, this results in divergent selection of mating-trait values between the two populations (Servedio, 2016). Hence, sexual selection mediated by (learning-based) assortative mating may promote hybrid zone stability in the form of geographic clines (Irwin, 2020; Brodin and Haas, 2009) or as mosaics, depending on the dispersal kernel (M’Gonigle and FitzJohn, 2010). These theoretical predictions are in accordance with empirical observations. Geographically structured polymorphism in phenotypes with relevance to mate choice are not uncommon in nature (McLean and Stuart-Fox, 2014), and learning of (self-) recognition cues or imprinting is widespread in the animal kingdom (Ten Cate and Vos, 1999; Bereczkei et al., 2004; Pfennig et al., 1983). Matching-rule-based mate choice thus features as a prevalent force in maintaining or promoting divergence with ongoing gene flow (Irwin and Price, 1999; Verzijden et al., 2012). This prediction applies particularly well to the context of hybrid zones in which mating traits that diverged in isolation come into secondary contact (Randler, 2008). Consistent with these predictions, the main model, best fitting the empirical data in the crow system, could explain the maintenance of steep clines over thousands of years for both loci associated with the mating trait (**Fig. 2**). However, all our models predicted that the hybrid zone will ultimately vanish, either as one color morph takes over, as in our best-fitting model, or as the clines will flatten out.

#### 4.1.1 Mode of mating preferences

The evolutionary outcome of sexual selection was contingent on the mating-preference function. Assuming self-reference for all possible seven phenotypes (**Fig. 1**) in the main model required little variation in mating probabilities (1.7-fold difference between minimal and maximal mating probabilities) to maintain high allelic differentiation between the parental populations. A different outcome was predicted for mate choice based on categorizing crows into three classes of pure parental and hybrid forms, as has previously been suggested for the crow system (Brodin and Haas, 2006, 2009). In the categorical model, the lack of discrimination among hybrid phenotypes increased gene flow of alleles from both loci with hybrids serving as a bridge for introgression between the pure parental forms (Irwin, 2020). Stronger avoidance (4.8-fold difference) of phenotypically dissimilar matings in the areas dominated by the parental genotypes was necessary to still obtain the empirically observed steep cline in genotypes and alleles of the mating-trait loci. This result demonstrates that the details of the preference function matter, even for a simple trait within a single sensory modality. This has potential implications for the evolution of matching-rule based mating preferences.

In the case of imprinting, many different modes are imaginable, including imprinting of hybrids to only one parental phenotype, to both, to intermediate phenotypes, or to the phenotypes of siblings or even other conspecifics of the local population. Imprinting to parental phenotypes may have similar effects as self-referencing as both result in positive frequency-dependent selection on the mating trait (Kopp et al., 2018). Yet, in simulations addressing this question we observed that cline dynamics depended strongly on the precise assumptions of imprinting (section 3.6). For the model with imprinting on one parent, cline dynamics mirrored the predictions from our main model with a simple self-referent matching rule. Simulations in which females imprinted on the mid-parent phenotype led to substantially different cline dynamics. Models explicitly addressing the interplay between the mating trait and the optimal resolution of preference for the trait seem to be worthy of further exploration. However, investigation of the empirical details underlying the matching rule (self-reference or imprinting and respective imprinting sets) and the actual mating cues would require large-scale cross-fostering experiments in the crow system.

#### 4.1.2 Genetic architecture of the mating trait

Hybrid-zone dynamics were strongly contingent on the genetic architecture of the mating trait. Gene flow was most readily reduced under an epistatic architecture, which fit the data best. Assuming an additive genetic architecture, we found no parameter combination that would recover cline steepness and offset of cline centers characteristic for the empirical data. Instead, under the best-fitting parameter values, the parental populations were rapidly homogenized by gene flow as illustrated by flattening of the clines across large geographic distances. Brodin and Haas (2009) modeled the southern-Danish hybrid zone between carrion crows in the south and hooded crows in the north. They assumed an additive model of three unlinked loci for the genetic basis of the mating trait without any other selection than assortative-mating-induced sexual selection. Assuming ongoing influx of hooded crows from the north and carrion crows from the south, their simulation runs led to stable phenotypic clines that resemble the cline observed in the field. However, with additivity of the mating trait it generally appears to be difficult to maintain divergence, as is illustrated by simulations from Irwin (2020) where very strong assortative mating was required for hybrid zone maintenance (10-fold difference). In these simulations, an increasing number of loci required even stronger assortment. These findings highlight the importance of the genetic architecture underlying the mating trait. Tentatively, these results may suggest that matching rules may be most relevant for evolution where single sensory modalities with a simple, at most oligogenic architecture (such as color (Cuthill et al., 2017)), are recruited for mate choice (Nadeau et al., 2007). Whether epistasis generally promotes divergence remains an open question and warrants further theoretical exploration. On the empirical side, genetic investigation of mating traits in systems where preference learning has been shown to play an important role for prezygotic isolation provides a promising empirical avenue (Yang et al., 2019). This likewise applies to sexually selected traits that appear to accelerate speciation rates on macroevolutionary scales (Panhuis et al., 2001; Hugall and Stuart-Fox, 2012; Seddon et al., 2013).

The genetic architecture also had a major effect on the spatio–temporal dynamics of the hybrid zone. Hybrid-zone movement is traditionally expected to be induced by dominance or constant asymmetries in allelic fitness, usually neglecting frequency dependence (Brodin et al., 2013; Mallet, 1986; Secondi et al., 2006) (for a mock exploration of our data with the model proposed by Barton, 1979, see online supplement section H.1). The effect of epistatis on hybrid zone movement has rarely been explored, least in the context of sexual selection. In this study, we observed that cline centers remained at the initial line of contact in the additive model, but shifted in all models when introducing an (empirically motivated, Knief et al., 2019) epistatic architecture. The combination of epistasis in the mating trait and frequency-dependent fitness effects had interesting emergent properties. The best-fit main model predicted an eastward movement of the hybrid zone which reversed its direction after ∼ 1,200 generations. This shift was due to changes in the relative allele frequencies of the two interacting loci. The Central European crow hybrid zone was mapped for the first time with great precision in 1928 (Meise, 1928). Re-examination of its location by Haas and Brodin (2005) 80 years (or ∼ 13 generations) later suggested a slight movement toward the east. While empirical sampling variance and sensitivity to model assumptions preclude a one-to-one comparison, both our results and those of Haas and Brodin predict that the crow hybrid zone does not conform to an equilibrium or quasi-equilibrium state and instead is dynamic to the present day. According to our model predictions, this will ultimately lead to the extinction of one or the other color morph within the time-frame of a glacial cycle.

#### 4.1.3 Releasing sexual selection

Last, we considered the effect of assortative mating alone without any induced fitness effects. A possible interpretation of this model is that there is a global, but no local carrying capacity, breeding sites are not limiting, and the time to find mating partners has no fitness effect. As a consequence, hybrid back-crosses with a phenotype resembling the pure morphs tend to leave the hybrid zone. Even though this model is entirely neutral, it allows for migration of individuals in search for appropriate mating partners. Accordingly, the model predicted a lower population density in the hybrid zone center and higher densities toward the parental ranges. While the model overall poorly supported the empirical geographic distribution of genotypes, it has interesting conceptual implications. In the popular tension-zone model, where hybrid zones are maintained at an equilibrium of migration and selection against hybrids, population density is preset by the local carrying capacity (Barton and Hewitt, 1985). Moving tension zones are predicted to be trapped in regions of low carrying capacity or biogeographic barriers to migration (Barton, 1979; Barton and Hewitt, 1985). On the contrary, our model predicts variation in population density as an emergent property of assortative mating in the absence of any form of selection. Hence, reduced population densities, as observed or predicted for hybrid zones (Barton and Hewitt, 1981, 1985; Hewitt, 1988), need not necessarily result from ecological constraints, but may be an emergent property of mate choice, even in the absence of selection.

### 4.2 Relevance to speciation

The genic view of speciation emphasizes that reproductive isolation will initially be caused by a small number of loci (Wu, 2001; Wolf et al., 2010; Presgraves, 2010). Divergent or disruptive selection reduces effective migration of alleles at these loci allowing for local allelic differentiation (Ravinet et al., 2017; Wolf and Ellegren, 2017). The mating-trait loci considered above constitute such barrier loci subject to selection (direct barrier effects). However, with few barrier loci of moderate effect, reproductive isolation remains confined to the linked local genomic neighborhood. Gene flow of neutral genetic variants disassociated from the selected background by recombination remains high. Indirect barrier effects reducing genome-wide gene flow are only expected to unfold when a sufficient number of barrier loci come into linkage disequilibrium—depending on the strength of selection and the recombination rate (Barton, 1983; Bierne et al., 2011; Feder et al., 2014). Here, we explored this indirect barrier effect by introducing a third, unlinked locus serving as a proxy for genome-wide, neutral genetic variation. Adding this locus to the best-fit main model allowed us to quantify genome-wide reduction in gene flow elicited by frequency-dependent sexual selection on a mating trait. While gene flow was slightly reduced, the effect was overall very small. This is consistent with theoretical models predicting that the effect of selected loci on neutral loci is small if the rate of recombination between these loci is large relative to the selection strength (Barton, 1986; Barton and Gale, 1993; Barton and Shpak, 2000b; Durrett et al., 2000; Sedghifar et al., 2016). Likewise, simulations by Irwin (2020), found that assortative mating had only minor effects on unlinked, neutral loci.

Speciation can be conceptualized as the built up of linkage disequilibrium between loci subject to divergent selection (Felsenstein, 1981). Eventually, this requires coupling of alleles across many barrier loci contributing to reproductive isolation (Wolf and Ellegren, 2017; Ravinet et al., 2017). In the crow system, however, it seems that only two polymorphic loci are associated with the mating trait and if there are no other loci under selection in the hybrid zone, only few genes may be affected by the hybrid zone. Indeed, Poelstra et al. (2014) found that only a small number of narrow genomic islands related to the plumage phenotype exhibited resistance to gene flow. The remainder of the genome appears not to experience any barriers to gene flow (see also Vijay et al., 2016). More specifically, Knief et al. (2019) demonstrated divergent selection in the European hybrid zone only for a ∼ 2 Mb region on chromosome 18 and the *NDP* gene (locus1 & 2, this study). Geographic clines were narrow for these loci (**see Fig. 2**), but stretched across several hundred kilometers for the remaining loci in the genome. Overall, these results are consistent with the findings of this study and suggest a plausible history of genome-wide admixture with the exception of few mating-trait loci maintained by preference learning. Divergence in putative mating trait (and preference) loci against a homogeneous genome-wide background is not restricted to the crow system (Toews et al., 2016; Hench et al., 2019; Malinsky et al., 2015; Wang et al., 2020). The framework presented here thus likely applies more widely, and makes it worthwhile exploring in other organisms tailored to the specifics of each system.

An important element of speciation is reinforcement of assortative mating (Yeh et al., 2018). If hybrids have reduced fitness, even if only in the sense of sexual selection induced by assortative mating, there may be selection in favor of mating strategies that avoid producing hybrids (Servedio and Kirkpatrick, 1997; Felsenstein, 1981). This reinforcement of assortative mating might be an important factor for speciation (Barton, 2013; Butlin and Smadja, 2018). It would be interesting to include reinforcement in our theoretical model to analyze how reinforcement could affect the dynamics of the hybrid zone and whether it could even lead to speciation before one of the two color morphs becomes fixed. The extent of reinforcement of assortative mating as an evolutionary process, however, is contingent on the amount of variation in the strength of mating preferences in the crow populations and on the genomic architecture of mating-preference behavior. To the best of our knowledge, both are unknown for crows, such that adding reinforcement to our theoretical model would be purely speculative or require extensive behavioral observations and population genomic analyses. As our model neglects the possibility of reinforcement and other eco-evolutionary aspects of the hybrid zone, we cannot rule out the possibility that speciation in carrion and hooded crows takes place. With our analyses, however, we show that the available data can also be explained by a model that will not lead to speciation.

## 5 Acknowledgments

We thank the members of the Wolf lab and the Dobzhansky discussion group at LMU Munich for valuable feed-back. For funding we thank the German Science Foundation DFG (grant ME 3134/6-2 to DM in Priority Program SPP 1590 “Probabilistc Structures in Evolution”), the European Research Council (ERCStG-336536 FuncSpecGen to JBWW) and LMU Munich (startup grant to JBWW).

