## Supplementary material for "Assortative mate choice and epistatic mating-trait architecture induce complex movement of the crow hybrid zone": Online Supplement

Dirk Metzler, Ulrich Knief, Joshua Penalba, Jochen B. W. Wolf
http://evol.bio.lmu.de/index.html

June 2, 2021

#### A Fitting model components to datasets

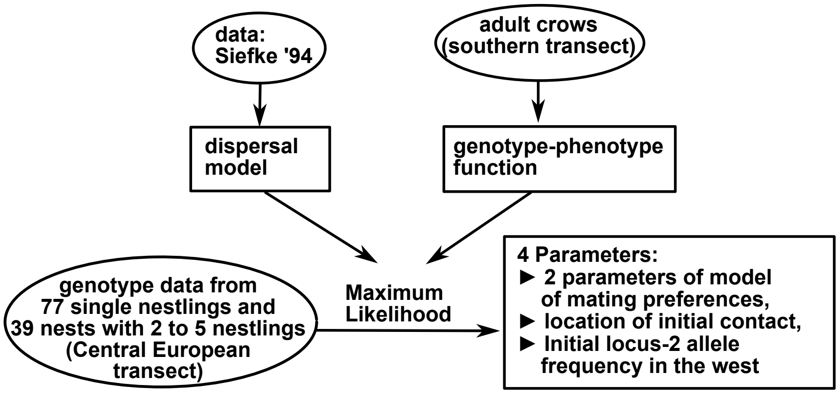


Figure S1: Overview of how different sources of data (ellipses) are used to fit the components of our model (boxes).

#### B Genotype–phenotype map

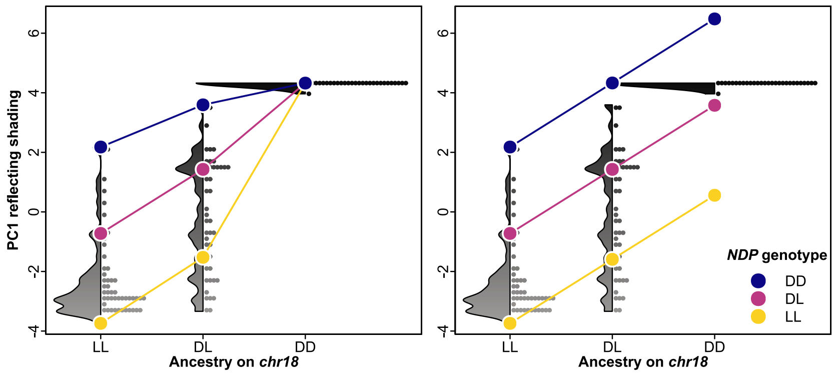


Figure S2: Phenotype estimates used in the simulation for an epistatic (left) and additive model (right).

### C Sampled genotypes

Figure S3 shows the genotype frequencies observed in the sampled chicks along the transect. Note that the data shown in most of the bars in figure S3 are not independent samples as multiple chicks were sampled from several nests. Sampling locations were binned into 5-km-bins and the km-numbers given in the figure are median values.

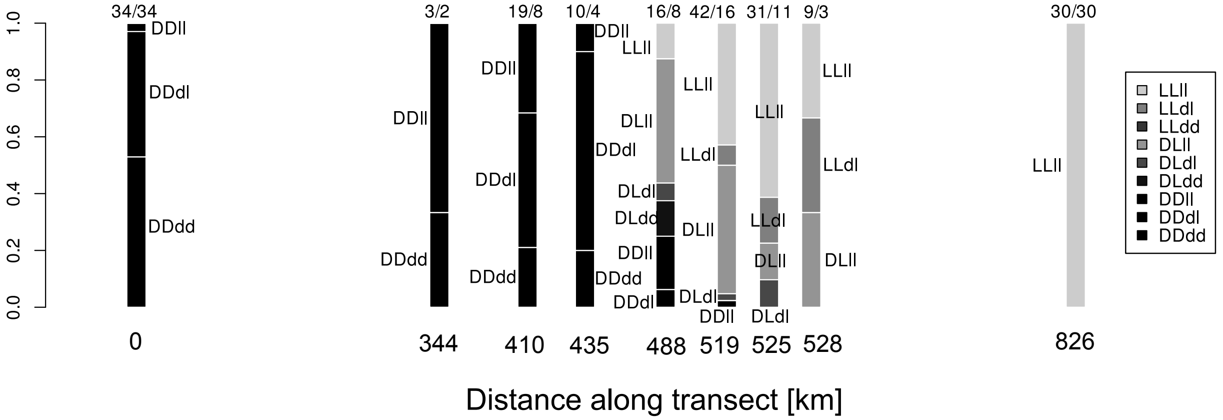


Figure S3: Observed genotype frequencies in chicks sampled along the Central European transect. The numbers above the bars are the sample sizes (number of crow chicks / number of nests from which they were sampled). Note that the genotype LLdd was not observed in these samples.

### D Dispersal

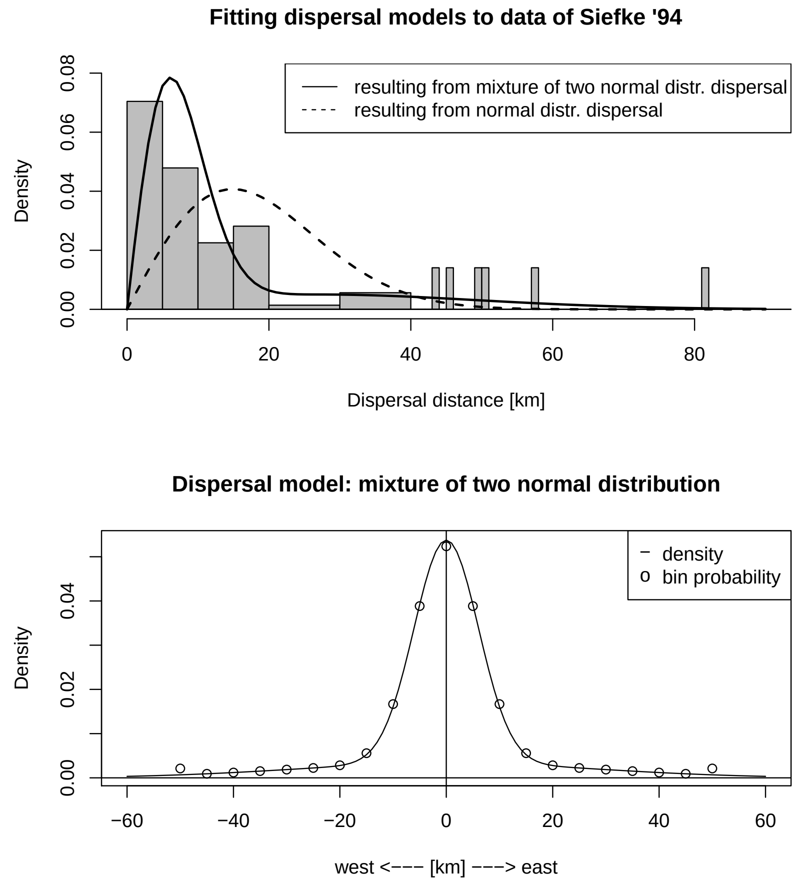


Figure S4: Fitting the Gaussian-mixture dispersal model to data of Siefke (1994). Distribution of dispersal distances in two dimensions (top) and the modeled distribution of dispersal orthogonal to the initial contact line (bottom).

#### E Mating-preference function

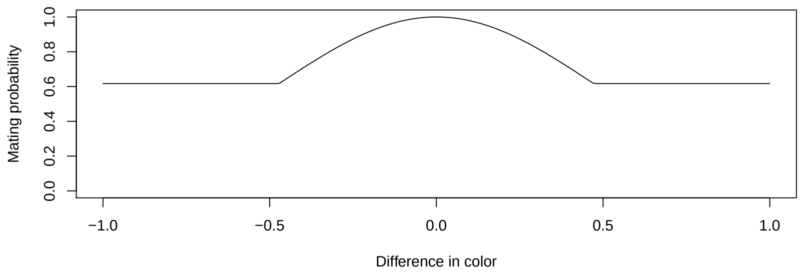


Figure S5: Mating preference $w_{g_1,g_2}$ as a function of phenotypic difference $\varphi(g_1) - \varphi(g_2)$, with parameter values $(\zeta, \eta) = (2.17, 0.617)$.

The resulting mating-preference values $w_{g_1,g_2}$ for all genotype combinations $(g_1, g_2)$ are shown in the main text in Fig.2B (top right).

#### F Alternative model without any fitness differences

Here we assume that all crows have the same mating chances, assuming that crows that find fewer mating partners compensate this by searching more intensely in their dispersal range, and we assume that this more intense search does not entail any fitness costs. The frequency of matings in bin $z$ between an individual of genotype $g$ stemming from bin $x$ and an individual $g'$ from bin $x'$ is then $w_{g,g'} \cdot f_{x,g} \cdot d_{x,z} \cdot v_{x,g} \cdot f_{x',g'} \cdot d_{x',z} \cdot v_{x',g'}$, where $w_{g,g'} \in [0, 1]$ is the mating attraction between individuals of genotypes $g$ and $g'$, which is 1 if $g$ and $g'$ induce the same phenotype. The factor $v_{x,g}$ makes sure that all individuals have the same fitness. The smaller the frequency of preferred mating partners of an individual $g$ in and around bin $x$, the larger is $v_{x,g}$. As this version of the model is supposed to be free of selection, we do not apply bin-wise rescaling as this would induce density-dependent competition and thus fitness differences.

In the simulation program we must numerically find the vector of values of $v_{\cdot,\cdot}$ that fulfill the following equations for all pairs $(x, g)$:

$$f_{x,g} = \sum_{z,x',g'} w_{g,g'} \cdot f_{x,g} \cdot d_{x,z} \cdot f_{x',g'} \cdot d_{x',z} \cdot v_{x,g} \cdot v_{x',g'}$$

or, equivalently,

$$1 = v_{x,g} \cdot \sum_{x'} \left( \sum_{z} d_{x,z} \cdot d_{x',z} \right) \cdot \sum_{g'} w_{g,g'} \cdot f_{x',g'} \cdot v_{x',g'}.$$

To apply a multidimensional Newton method available in the GNU Scientific Library (GSL) to compute the values of $v_{x,g}$, we need some derivatives. The derivative of the right-hand side with respect to $v_{x,g}$ is:

$$\sum_{x'} \left( \sum_{z} d_{x,z} \cdot d_{x',z} \right) \cdot \sum_{g'} \left(1 + I_{(x',g')=(x,g)}\right) \cdot w_{g,g'} \cdot f_{x',g'} \cdot v_{x',g'}$$

The derivative with respect to any other $v_{x',g'}$ (that is $x' \neq x$ or $g' \neq g$ or both) is:

$$v_{x,g} \cdot \left( \sum_{z} d_{x,z} \cdot d_{x',z} \right) \cdot w_{g,g'} \cdot f_{x',g'}$$

The sums over $z$ are set into parentheses here just to indicate the following: To accelerate the numerical solution we store the values $\sum_z d_{x,z} \cdot d_{x',z}$ in a 2-dimensional array of size $200 \times 41$, where the second index is $x' - x + 20$.

### G Results

#### G.1 Best fitting model (with induced sexual selection)

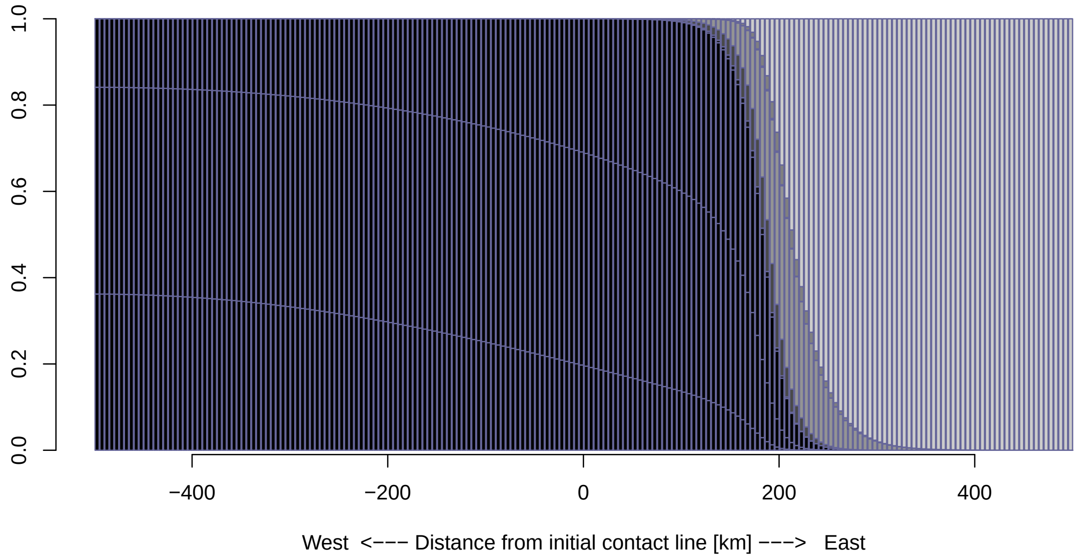


Figure S6: Genotype frequencies in each bin after 2000 generations in the main model. From bottom-left to top-right: *DDdd*, *DDdl*, *DDll*, *DLdd* (very rare), *DLdl*, *DLll*, *LLdd* (very rare, hard to see in Figure), *LLdl*, *LLll*. Grey tones mimic average grey tone of corresponding crow phenotypes.

#### G.2 Neutral model

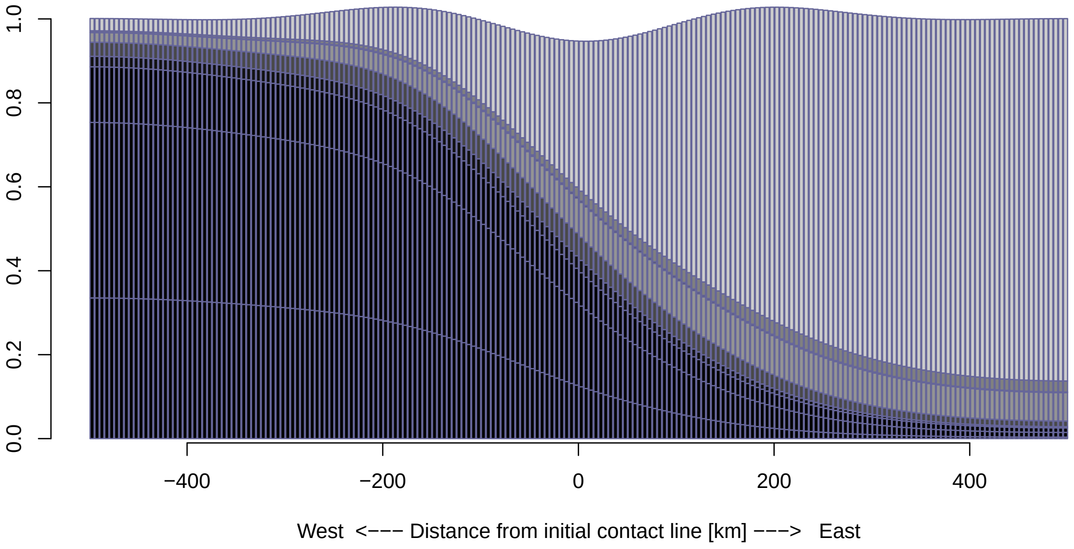


Figure S7: Genotype frequencies in the neutral model after 2000 generations. From bottom-left to top-right: DDdd, DDdl, DDll, DLdd, DLdl, DLll, LLdd (very rare, almost invisible in Figure), LLdl, LLll. Grey tones mimick average grey tone of corresponding crow phenotypes. For the scale on the vertical axis: initial cumulative genotype frequency was 1.0 in each bin.

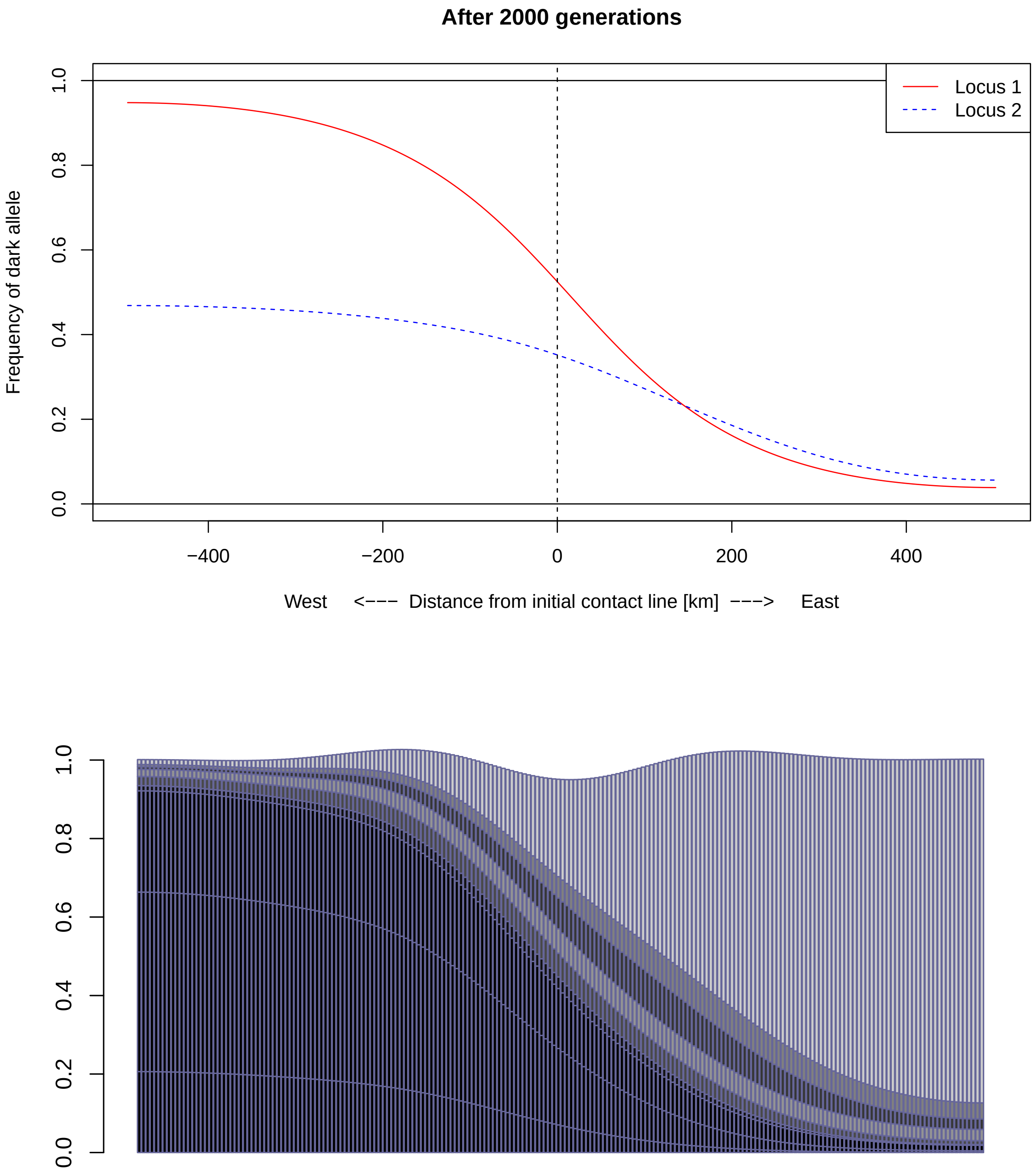


Figure S8: Results from neutral model with $(\zeta, \eta) = (300, 0.05)$ and an initial locus 2 dark allele frequency of 0.6. From bottom-left to top-right: DDdd, DDdl, DDll, DLdd, DLdl, DLll, LLdd, LLdl, LLll.

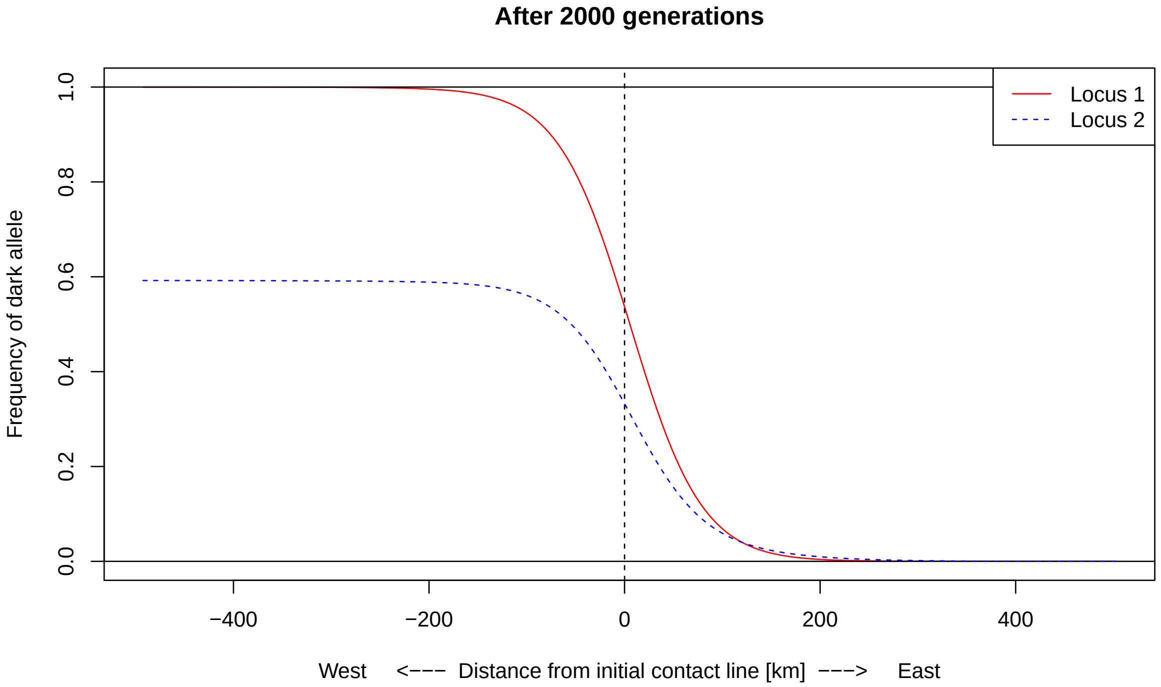


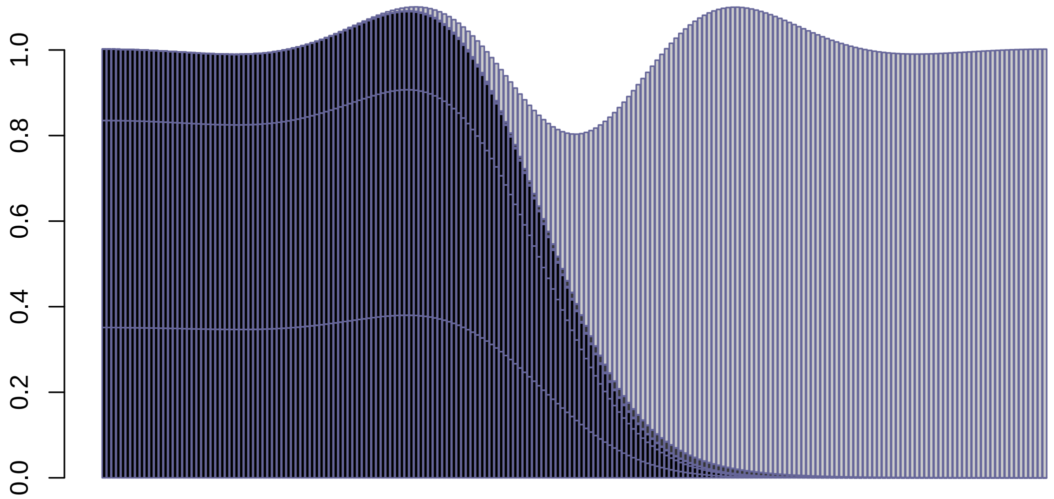


Figure S9: Results from neutral model with $(\zeta, \eta) = (300, 0.001)$ and an initial locus 2 dark allele frequency of 0.6. The only hybrid genotype that reaches frequencies above 0.00125 is LLdd, whose frequency is slightly above 0.019 in the center of the hybrid zone.

#### G.3 Assuming an additive genomic architecture of coat color

When we tried a parameter combination with more mating assortativity ($\rho = 2.7, \eta = 0.6$) than in the best fitting additive model, we observed that the clines for the two loci clearly differed after 2000 generations, but we did not observe the east–west displacement of the clines that we see in the empirical data (Figure S10).

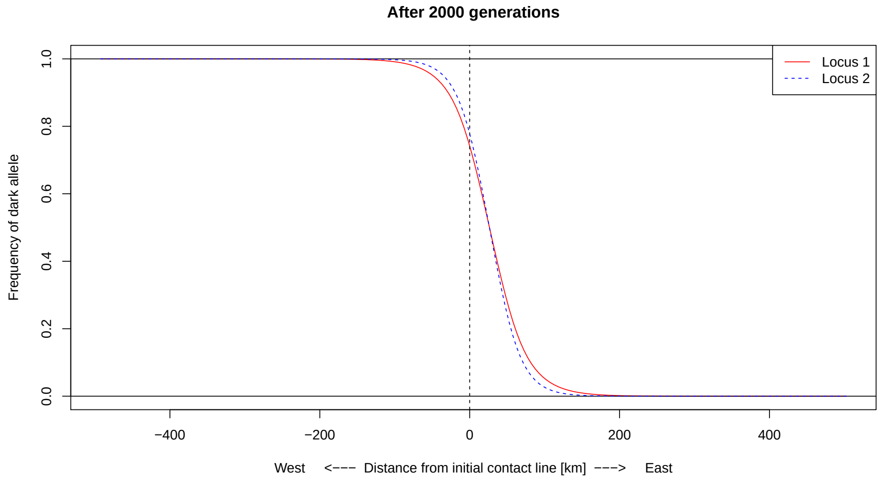


Figure S10: Results from model with an additive genetic architecture of the coat color trait, initially fixed alleles in the west and in the east and parameter values $(\zeta, \eta) = (2.7, 0.6)$.

When we allowed that locus 2 was initially polymorphic in the west (which means that not all crows were black there), we obtain maximum-likelihood parameters of $\rho = 1.832$, $\eta = 0.967$, and 0.56 for the initial allele d frequency in the west at locus 2. In this scenario, the frequency of allele D at locus 1 leveled out to a value far below 1 in the west (Figure S11).

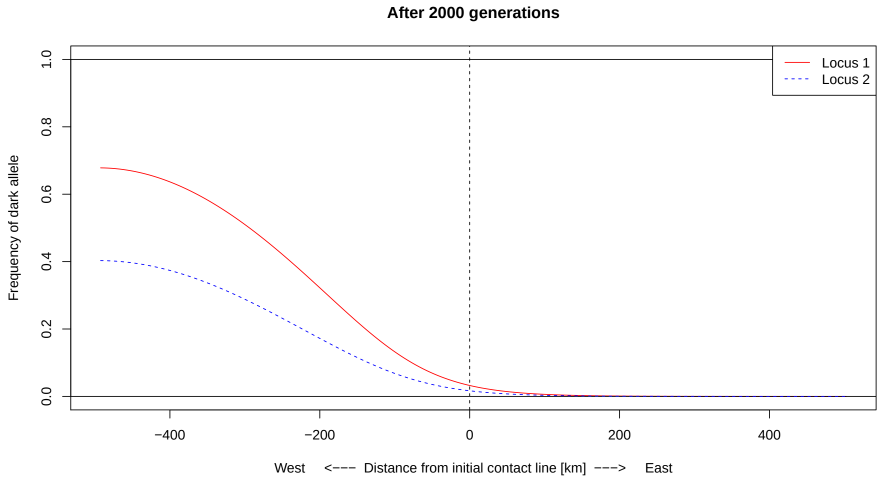


Figure S11: Results from model with an additive genetic architecture of the coat color trait, initially allele d frequency of 0.56 in the west and in the east and parameter values $(\zeta, \eta) = (1.832, 0.967)$. (These parameter values were obtained by likelihood optimization.)

#### H Shift of hybrid zones

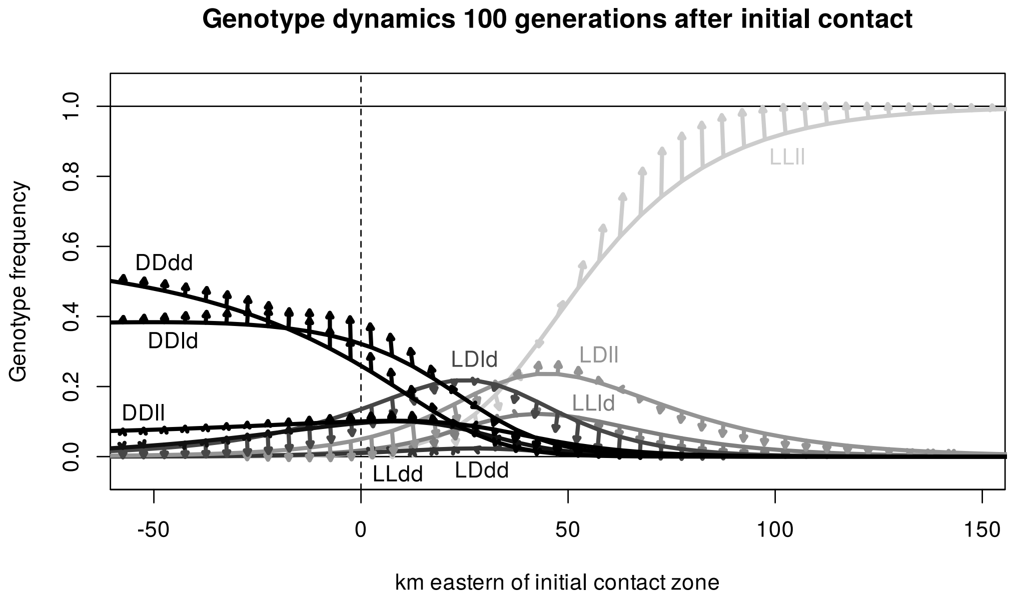


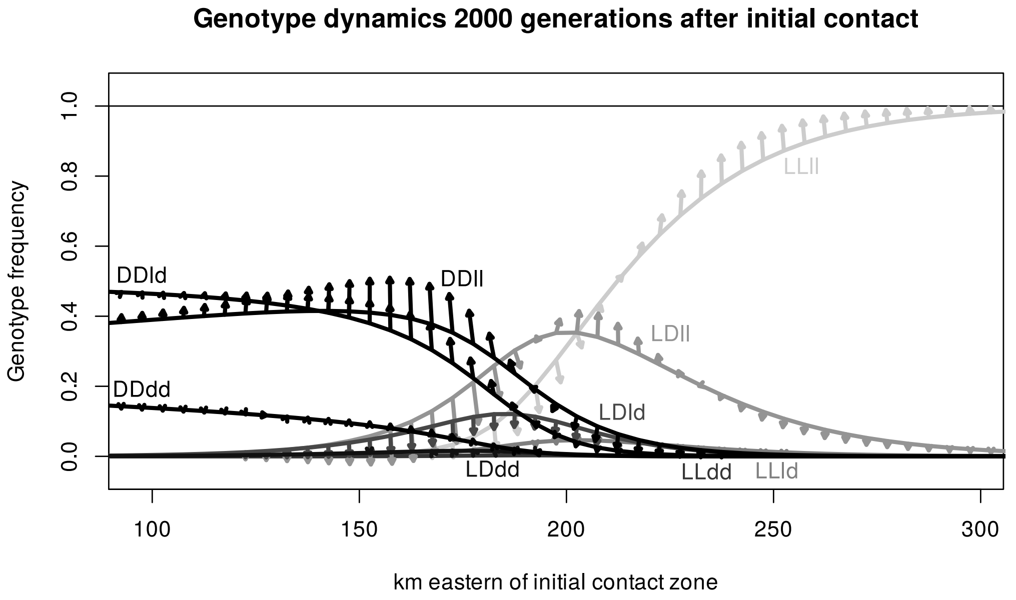


Figure S12: Genotype frequencies around the hybrid zone after 100 and after 2000 generations after initial contact. Arrows pointing up or down indicate that carriers of the genotype produce offspring number that is above or below average, respectively. The horizontal component if tilted arrows the average direction in which carriers of the genotype at that position find mating partners and breed.

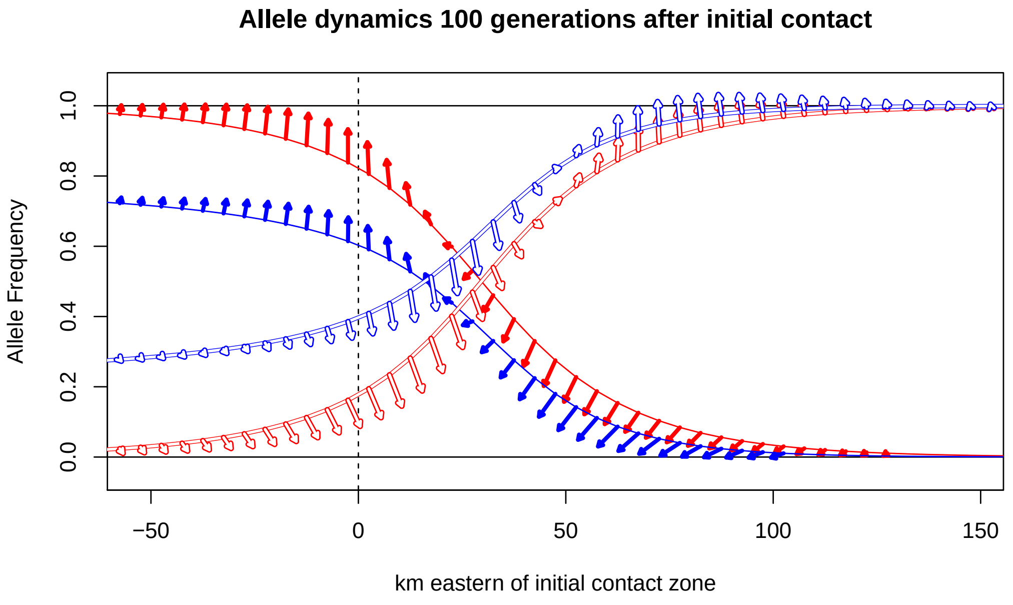


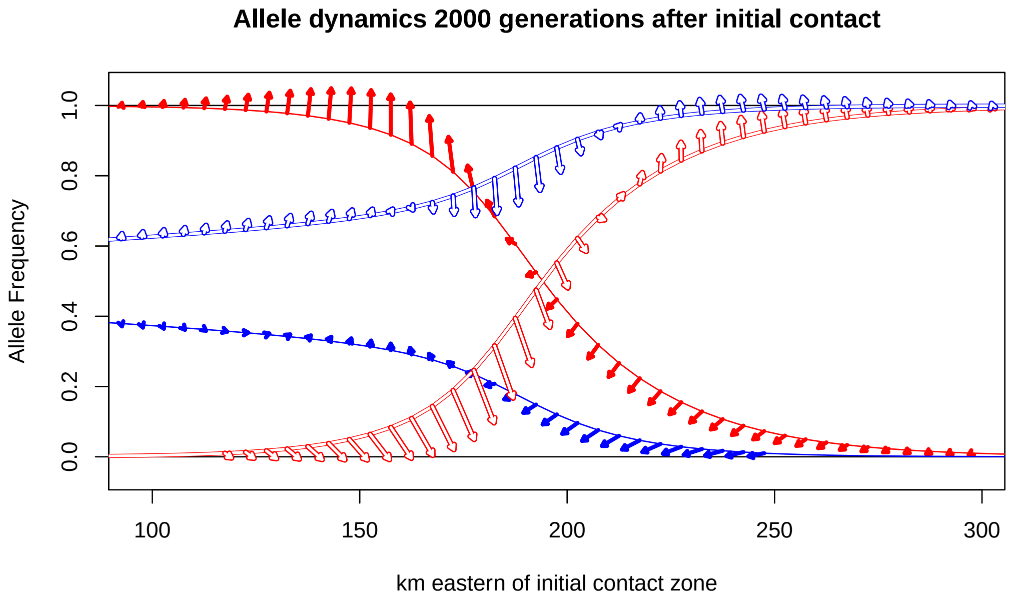


Figure S13: Allele frequencies at locus 1 (red) and locus 2 (blue) around the hybrid zone after 100 and after 2000 generations after initial contact. Solid lines represent the "dark" alleles $D$ and $d$, and white lines with red or blue margin represent the "light" alleles $L$ and $l$. Arrows pointing up or down indicate that carriers of the allele produce offspring number that is above or below average, respectively. The horizontal component if tilted arrows represent the average direction in which carriers of the allele at that position find mating partners and breed.

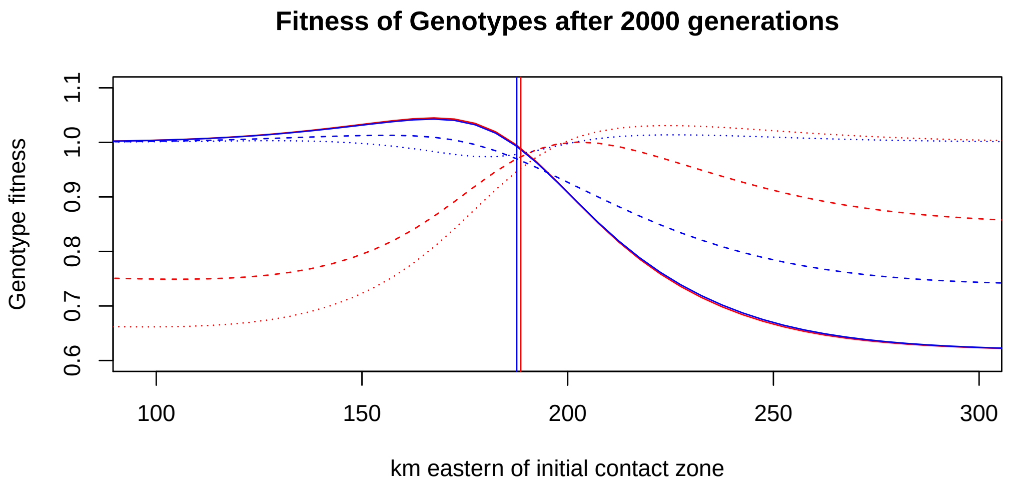


Figure S14: Fitness values of single-locus genotypes at locus 1 (red) and locus 2 (blue) around the hybrid 2000 generations after initial contact. Solid lines represent the "dark" genotypes $DD$ and $dd$, dashed lines the heterozygotes $DL$ and $dl$ and dotted lines for the "light" genotypes $LL$ and $ll$. Vertical lines show the centers of the clines of the two loci, that is the points of inflection, at that time point.

#### H.1 Comparison to a classical prediction of cline movement

Barton (1979) proposed a hybrid zone model in which the speed of the movement of clines can be calculated from dispersal and selection parameters. The model assumes natural selection with fitness values of genotypes that do not vary in space. Even though this assumption does not apply to our scenario where the fitness of genotypes is frequency dependent (Figure S14), we can still explore whether we can Barton's model to explain the movement of the cline centers.

For locus 1 the prediction of our best-fitting model for the present time, that is 2000 generations after initial contact, is that the average offspring numbers of individuals with genotypes $DD$, $DL$ or $LL$ that stem from the center of the cline are 0.989, 0.973, and 0.952, respectively (Figure S14) . To scale these fitness values and express them like in Barton (1979) as $1 + 2\alpha s$, $1 + (\alpha - 1)s$ and 1 we have to set $s = -0.00333$ and $\alpha = -5.76$. This, however, means that we are not in the scope of the model in Barton (1979), which assumes positive values for $s$ and $\alpha$.

For locus 2 the average offspring numbers of genotypes $dd$, $dl$ and $ll$ are 0.993, 0.97 and 0.978, corresponding to $s = 0.0158$ and $\alpha = 0.477$. In the model of Barton (1979), a cline moves with a speed of $\alpha \cdot \sqrt{s \cdot m/2}$ in the direction to where the less fit genotype is more frequent, where $m$ is the variance of the dispersal kernel. The one-dimensional dispersal kernel that we fitted to the data of Siefke (1994) has a variance of 190.58 km$^2$, which leads to $\alpha \cdot \sqrt{s \cdot m/2} \approx 0.585$ km/generation. As the genotype $dd$ produces on average more offspring than $ll$ and is more frequent in the west, the cline would move eastward if the cline behaved according to the model of Barton (1979). In our simulation we observe a westward movement of 0.587 km per generation for the time point 2000 generations after initial contact of the locus 2 cline, but the direction of movement ist westward.

If we caclulate the locus 2 fitness values not only for crows that stem from the center of the cline, but for a range from 20 km western to 20 km eastern of the locus 2 cline center, the average offspring numbers for $dd$, $dl$ and $ll$ are 0.964, 0.958 and 0.986, such that $ll$ has the highest fitness. With $s = 0.0182$ and 0.629 we can express the rescaled fitness values as 1, $1 + (\alpha - 1)s$, and $1 + 2\alpha \cdot s$, for which the model of Barton (1979) would predict a movement of the cline with 0.828 km per generation to the west. Of course, this value will be different if we choose a different range around the cline center or if we use the dispersal kernel conditioned on mating success, perhaps even stratified for the different genotypes. For locus 1 the genotypes fitness values in a range from 20 km western to 20 km eastern of the cline center still leads to negative values of $s$ and $\alpha$.

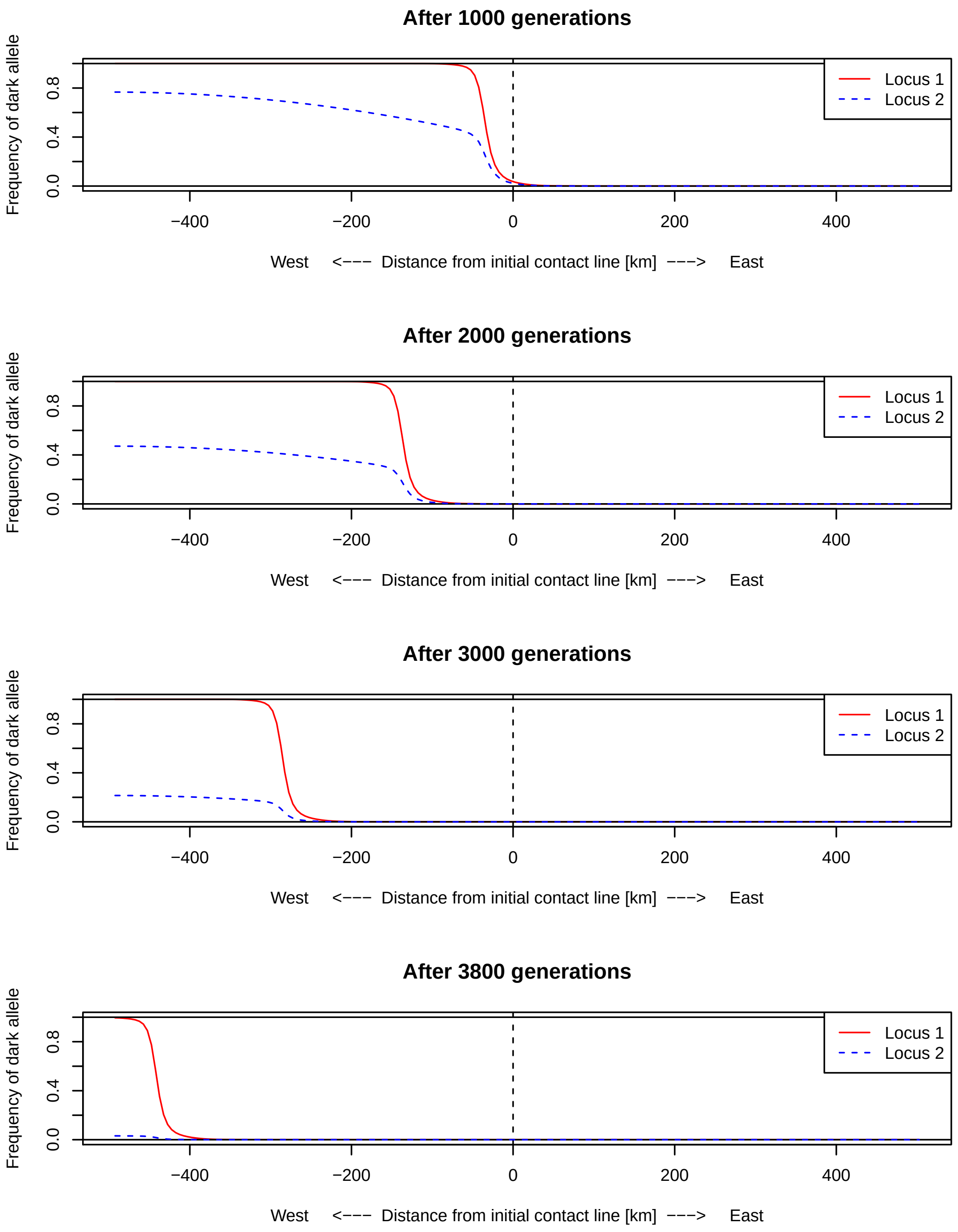


Figure S15: According to the categorical model, the hybrid zone first shifts westwards until the hooded crows go extinct.

### I Almost no effect on an unlinked neutral locus

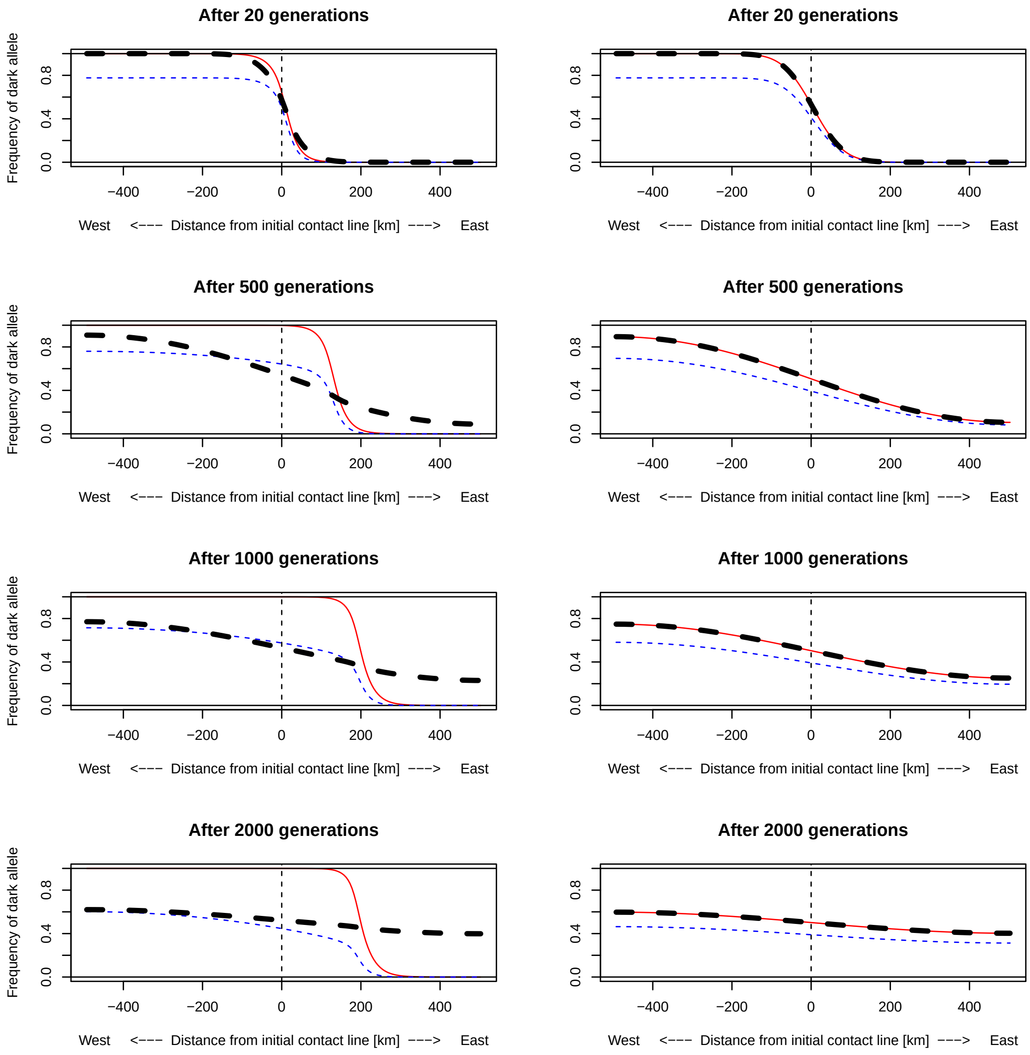


Figure S16: Left column: only the third locus (dashed bold line) is neutral, Right column: all three loci are neutral.

#### I.1 Movie of simulation results

A movie version of these simulation results is available on
`https://raw.githubusercontent.com/statgenlmu/assortative_crows/master/movies/threeloci_6000.mp4`.

The movie shows hybrid-zone movement through time shown for both mating-trait loci (locus 1, locus 2) and an unlinked, neutrally evolving third locus (locus 3). Allele frequencies are shown on the y-axis for the dark alleles of locus 1 (red line) and locus 2 (blue line) which define the genotype phenotype map (see Fig. 1 of the main manuscript). Allele frequency changes can be followed from the initiation of secondary contact at generation 0 over 6,000 generations (corresponding to 36,000 years) along a one-dimensional, spatial representation of the hybrid zone (x-axis). Upper panel: results from the main model invoking sexual selection. Lower panel: results from the same model assuming random mating precluding any form of assortment by phenotype.

### J Finite-population models

In this class of simulation models we assume that nesting places of crows are located on a 2-dimensional $1000 \times 500$ grid, where the grid points have a distance of 1 km to their neighbors and 1000 km is the expansion in west-east direction. In other words, each crow nest is assumed to be in the center of territory of 1 km$^2$. For the initial condition we assume that all crows in the $500 \times 500$ western nests have the carrion-crow genotype and all crows in the $500 \times 500$ eastern nests have the hooded-crow genotype. We assume discrete generations. In each generation we sample for each nest (that is, grid point) a pair of nests according to the dispersal model. For these two nests, offspring are generated accoring to the parents that occupied these nests in the previous generation. Based on the mating-preference model a decision is then made whether the two generated crows mate. If they mate, they occupy the focal nest. If they do not mate, the procedure is repeated (including the sampling of a pair of parental nests) until two crows mate and occupy the focal nest. The dispersal model according to which we choose the parental nests is based on the mixture of two conical two-dimensional normal distributions that we fitted to the data of Siefke (1994), see main text section 2.2.4. Like in the main model, the boundaries of our simulated space are reflecting, but in contrast to the main model, dispersal distance is not limited to 50 km in the finite-population model.

This modeling approach is more flexible and allows us to explore also certain imprinting models in addition to the finite-population-size variant of our main model with self-referencing. Fitting parameters with simulations directly based on the finite-population models would, however, be computationally very demanding, such that we only carry out simulations with parameter combinations that are based on our parameter estimations for the main model (see main text section 3.1).

#### J.1 Self-reference finite-size model

In this variant of the finite-population model the same self-reference mating preference function as in the main model is applied with the model parameters fitted for the main model (see main text section 3.1).

Figure S17 shows the spatial distribution of phenotypes from one simulation run at time points of 1, 2000 and 5000 generations after initial contact, and figure S18 shows the corresponding allele frequency clines along the transect. Like in our main model, the hybrid zone first moves eastward and then westward (compare to main-text figure 3). The same dynamics was observed in independent replications of the simulation (data not show).

#### J.2 Female-choice model with imprinting on mid-parental phenotype

We now assume that the fameles are the ones who make the mating decision, and it is based on the similarity of the candiate male to the average phenotype of the female's parents. The mating probability is still given by the mating preference function inferred in main text section 3.1, but this time applied to the difference between the phenotype of the candidate male and the average phenotype of the female's parents. (Note that there is not other difference between males and females in our models, such that we obtain an equivalent male-choice model if we simply exchange the roles and females in mate choice.)

Figure S19 shows the allele frequency clines after 1, 1000 and 2000 generations of one simulation run. Compared to our main model and of the finite-populations self-reference model, there were two qualitative differences in the allele-frequency cline dynamics that we observed in the simulations with the mid-parental imprinting model. First, the hybrid zone moved eastward until all crows were black and, second, at some time point (after 1000 generations) there was a range in the hybrid zone in which the dark-allele frequency at locus 2 was higher than the dark-allele frequency at locus 1. We observed this also in independent replications of the simulation (data not shown).

#### J.3 Female-choice model with imprinting on father (or mother)

Like in J.2 we assume female-choice but in this model variant the female compares the phenotype of the candidate male to the phenotype of her father, again applying the mating preference matrix inferred in main text section 3.1. In another sub-variant of the model, females are not imprinted to their fathers' phenotype but to those of their mothers. (Again we obtain equivalent male-choice models by swapping the roles of the sexes.)

Figure S20 shows that the allele cline dynamics observed in a simulation of this model was qualitatively equal to the dynamics oberverd in the simulations of our main model (main text figure 3) and of the finite-populations self-reference model (figure S18). With some quantitiative variation, this was also the case for independent replications of the simulations, also when imprinting to the mother-phenotype was assumed (figure S21).

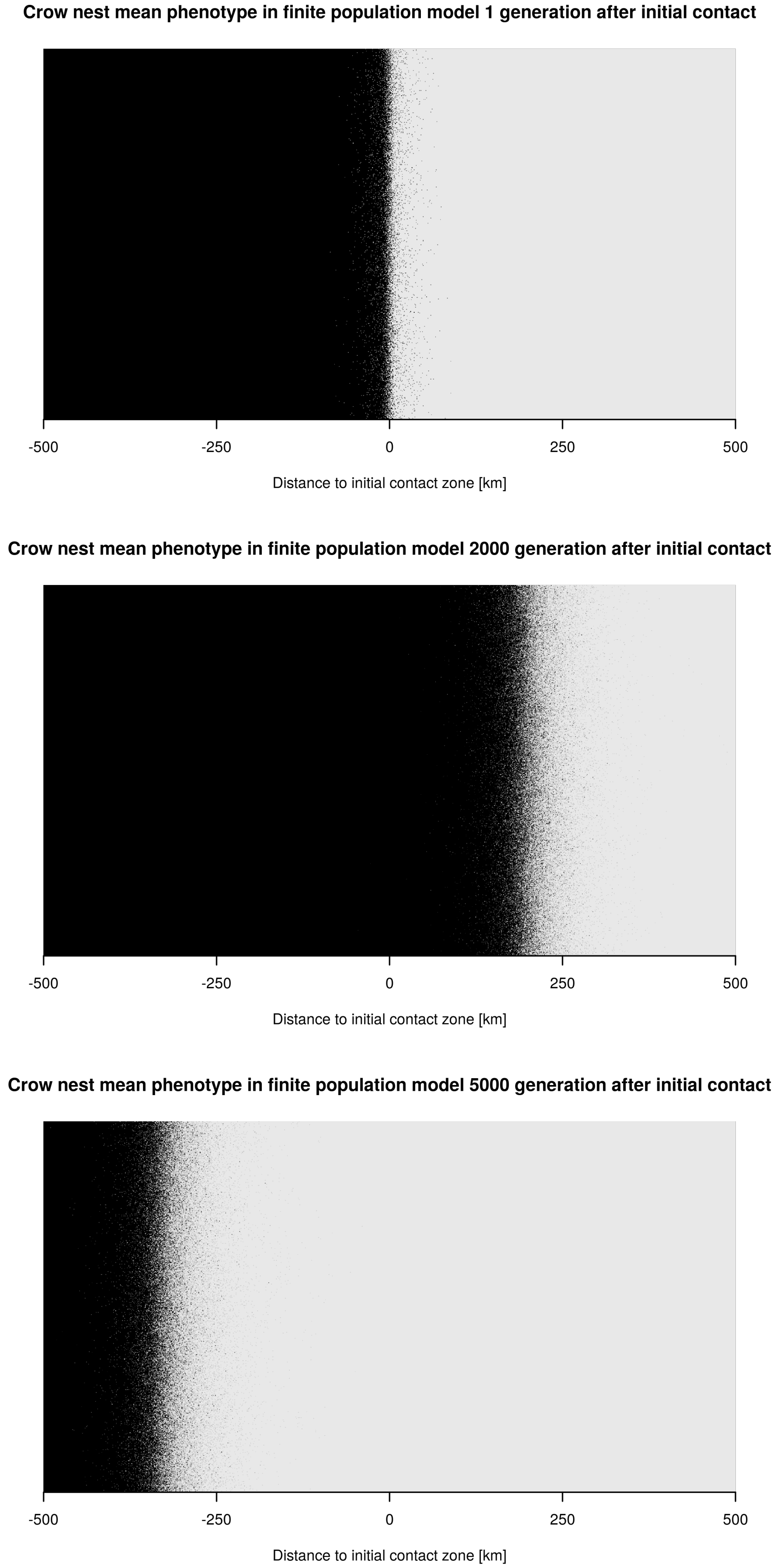


Figure S17: Phenotype distributions in a finite-population self-reference model simulation run after 1, 2000 and 5000 generations after intial contact. The grey tone of each pixel represents the mean phenotype of the parents in the nest.

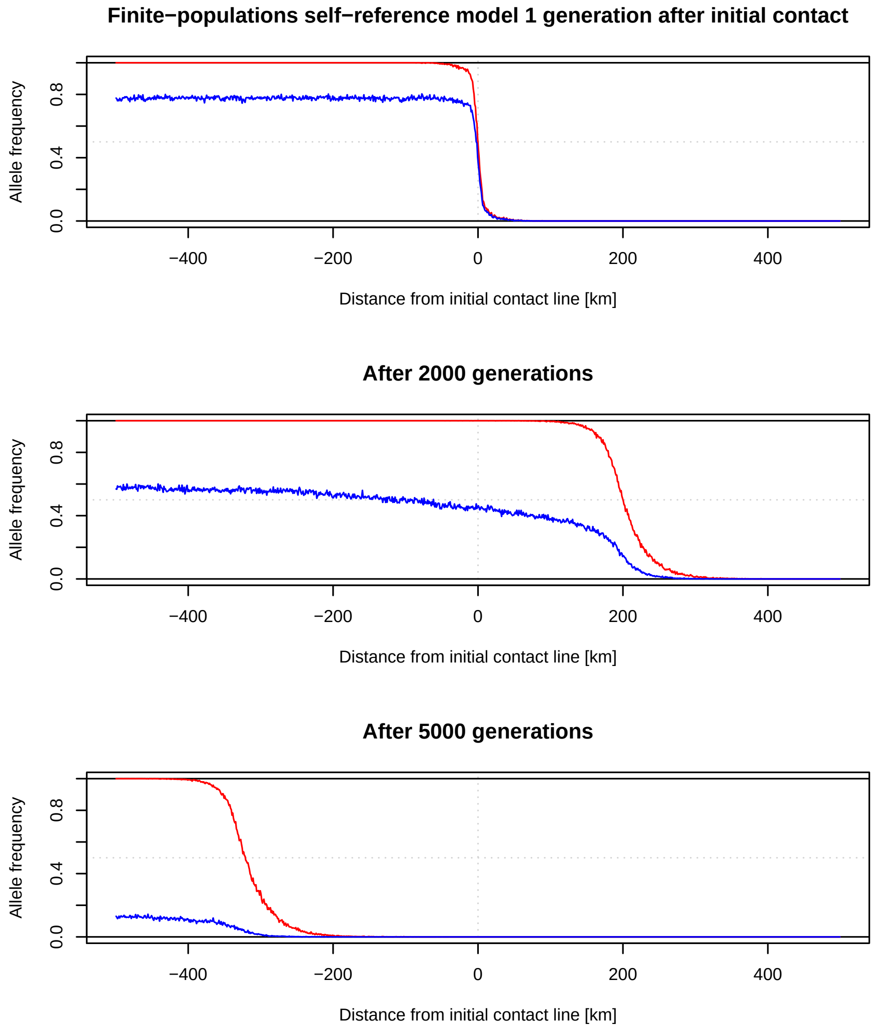


Figure S18: Clines of the alleles for darker colour for the locus 1 (red) and locus 2 (blue) after 1, 2000 and 5000 generations after initial contact according to a simulation run with the finite-population self-reference model.

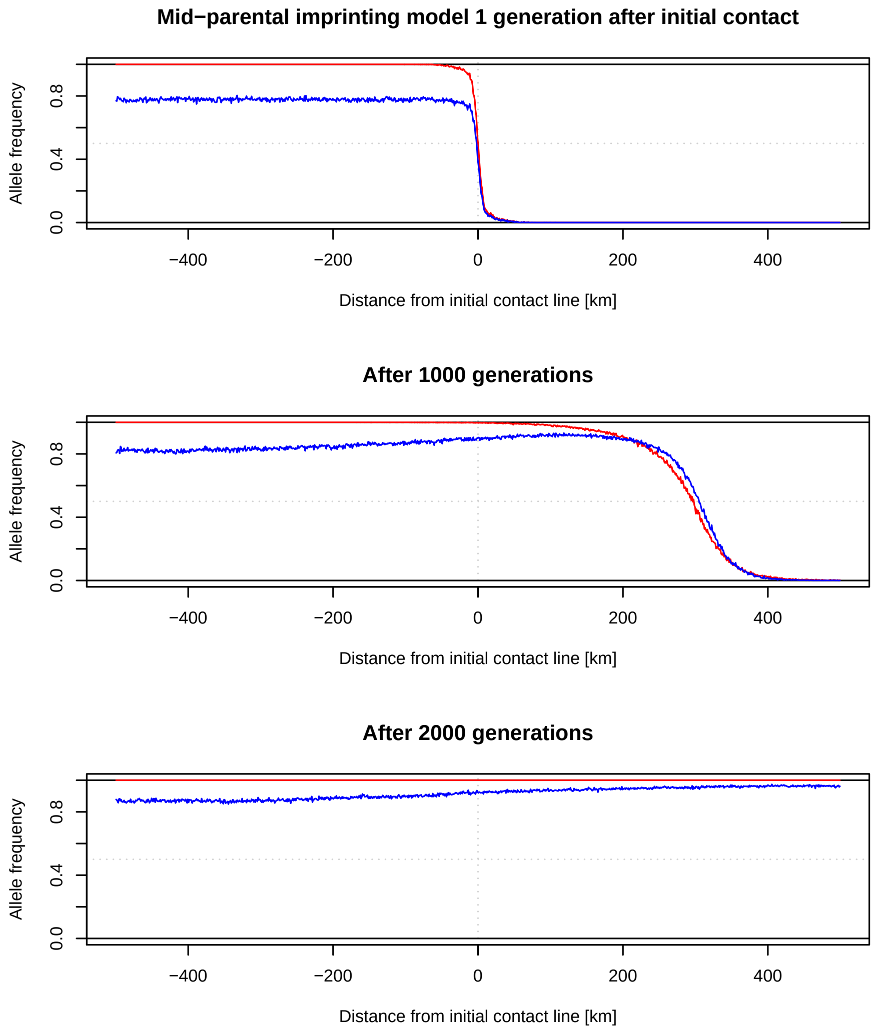


Figure S19: Clines of the alleles for darker colour for the locus 1 (red) and locus 2 (blue) after 1, 1000 and 2000 generations after initial contact according to a simulation run with the finite-population with female-choice and the imprinting of females according to the average phenotypes of their parents.

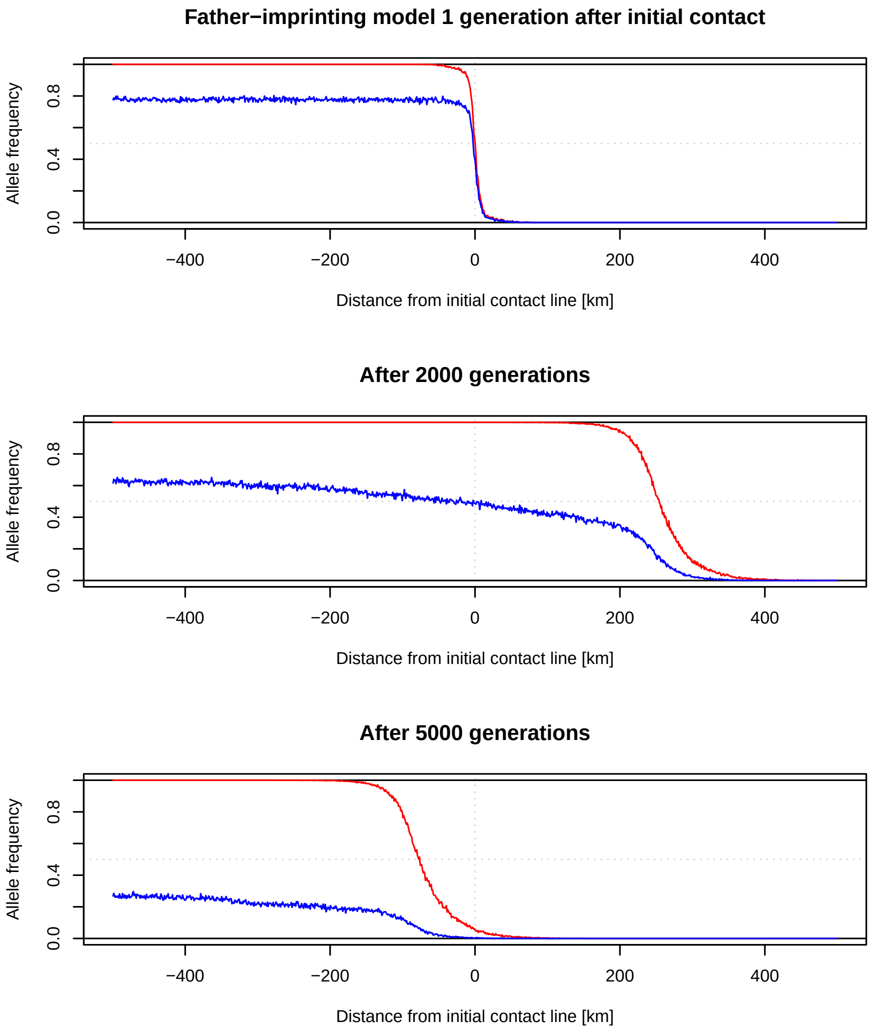


Figure S20: Clines of the alleles for darker colour for the locus 1 (red) and locus 2 (blue) after 1, 2000 and 5000 generations after initial contact according to a simulation run with the finite-population with female-choice and the imprinting of females according to the phenotypes of their father.

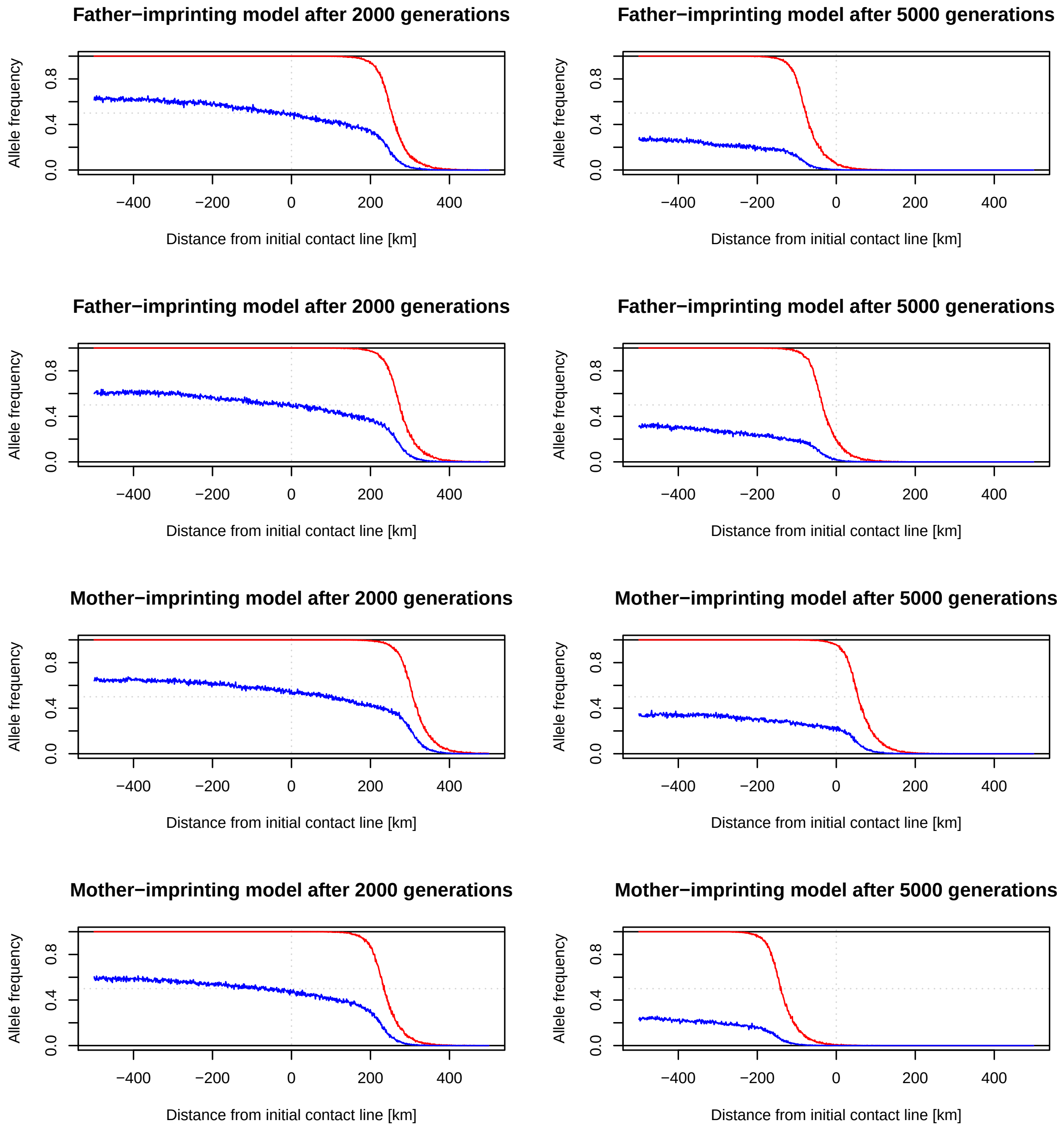


Figure S21: Clines of the alleles for darker colour for the locus 1 (red) and locus 2 (blue) after 2000 (left column) and 5000 generations (right column) after initial contact according to four independent simulation runs with the finite-population with female-choice and imprinting. Plots that are shown next to each other in the same row show data from the same simulation run. In two runs (two top rows) it was assumed that females were imprinted to the phenotypes of their fathers and in the other two (two bottom rows) it was assumed that they were imprinted to the phenotype of their mothers.
